# Anti-*Toxoplasma gondii* screening of eight species and bio-guided identification of metabolites of *Elaeis guineensis* leaves, a preliminary study

**DOI:** 10.1101/2024.06.24.600452

**Authors:** Nangouban Ouattara, Abdulmagid Alabdul Magid, Sandie Escotte-Binet, Philomène Akoua Yao-Kouassi, Isabelle Villena, Laurence Voutquenne-Nazabadioko

**Affiliations:** Université de Reims Champagne-Ardenne, CNRS, ICMR, Reims, France; Université de Reims Champagne Ardenne, ESCAPE, Reims, France; Centre National de Référence de la Toxoplasmose, CHU Reims, 51092 Reims, France; Laboratoire de Constitution et Réaction de la Matière, UFR Sciences des Structures de la Matière et de Technologie, Université Félix HOUPHOUET-BOIGNY 22 BP 582 Abidjan, Côte d’Ivoire; Université Polytechnique de San Pedro, BP V1800 San Pedro, Côte d’Ivoire

**Keywords:** *Toxoplasma gondii*, Vero cell, *Elaeis guineensis*, bioguided fractionation, dereplication, medicinal plants

## Abstract

The apicomplexan parasite *Toxoplasma gondii* causes toxoplasmosis, a ubiquitous and cosmopolitan parasitosis, generally asymptomatic and potentially dangerous for the fetus and highly immunocompromised patients. Pyrimethamine and sulfadiazine, supplemented with folic acid, are the drugs of choice to treating the disease, but they produce severe side effects and treatments fail due to drug resistance. New anti-*Toxoplasma* compounds are needed, and natural compounds can be a good source for obtaining them. The antiparasitic activity of 40 polar and non-polar extracts of eight antiparasitic medicinal plants used in Côte d’Ivoire, and selected based on ethnopharmacological survey, were evaluated in *vitro* against *T. gondii*. Among them, the hydromethanolic extract of the *Elaeis guineensis* leaves exhibited the best parasite growth inhibition (94% ± 0.07) at 25 μg/mL without being cytotoxic at the same dose. The fractionation of this extract did not allow the recovery of antitoxoplasmic activity in its individualized fractions. The ^13^C-NMR based dereplication of the produced fractions and the purification of one of them highlighted the presence of saccharides, terpenes, flavonoids, cardanols and aliphatic acids. The obtained clusters of ^13^C chemical shifts were assigned to their corresponding molecular structures with the help of open online databases, resulting in nine unambiguously identified compounds, whereas the purification of one of these fractions led to the identification of palmitic acid (**10**), palmitic acid methyl ester (**11**), 3-[12(*E*)-pentadecenyl]phenol (**12**), 3-[14(*E*)-heptadecenyl]phenol (**13**), ursolic acid (**14**) and stigmasta-4,24(28)-dien-3-one (**15**). A synergistic effect between these metabolites is thought to be responsible for the anti-*Toxoplasma* activity.

GRAPHICAL ABSTRACT

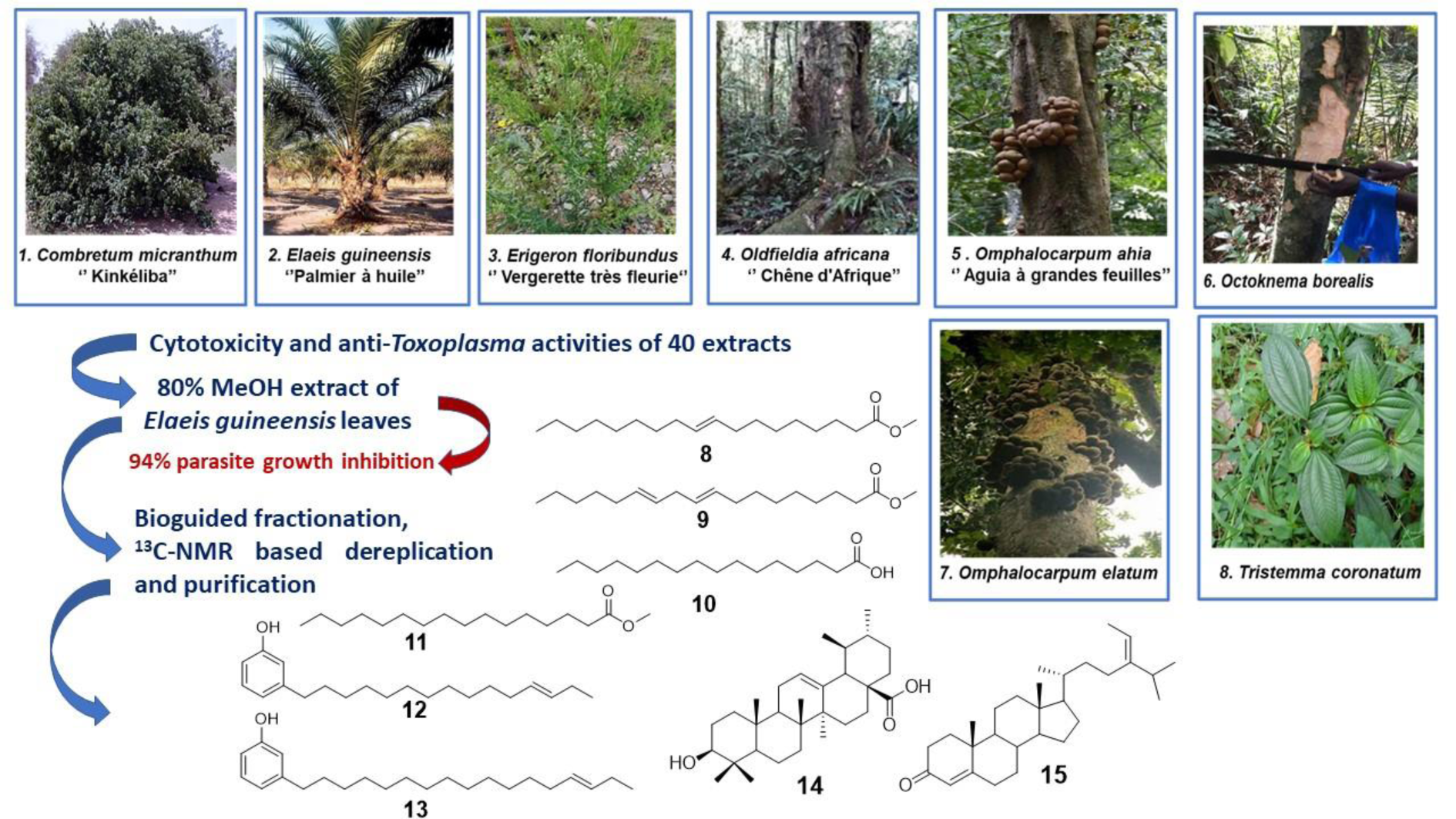

**Highlights:**

- Antiparasitic activity of 40 polar and non-polar extracts of eight plants were investigated
- The hydromethanolic extract of the *E. guineensis* leaves exhibited 94% parasite growth inhibition
- The fractionation of this extract did not allow the recovery of antitoxoplasmic activity
- Nine compounds were identified by ^13^C-NMR based dereplication
- Six compounds were purified from the cytotoxic fraction

---

Toxoplasmosis is a ubiquitous disease of parasitic origin caused by an obligate intracellular eukaryote *Toxoplasma gondii,* whose final host is a felid (1, 2). Although most human infections are uncomplicated, the parasite can persist and the infection can be reactivated in immunosuppressed patients leading to cerebral or generalized toxoplasmosis in immunocompromised patients (2). In add, when infection is acquired during pregnancy, it can cause congenital toxoplasmosis with serious consequences for the fetus or the infant (3−5). *T. gondii* could also cause severe ocular disease in immunocompetent patients (6, 7) and in congenital toxoplasmosis (8).

It is estimated that about 30-50% of the world’s population is infected with *T. gondii* (9). Toxoplasmosis is ranked as the third largest contributor to the health burden caused by foodborne illness in Europe (10). In France, prevalence of toxoplasmosis is decreasing and estimated around 31% in 2016 (11), with around 150-200 newborns affected by congenital toxoplasmosis each year. Similarly, in Côte d’Ivoire, the latest Epidemiological studies have shown that seroprevalence has decreased from 80% in the 1960s to 45 % in 2014 (12).

The current treatment of acute toxoplasmosis is the combination of sulfadiazine and pyrimethamine, which act synergistically to block the folate pathway, involved in DNA synthesis with the dihydrofolate reductase (DHFR) and dihydropteroate synthetase (DHPS) enzymes. Pyrimethamine act on parasite DHFR, but are unable to distinguish it from the enzyme of the human host, whereas sulfadiazine block DHPS, and leucovorin to minimize host bone marrow toxicity (13). Unfortunately, this inhibition also creates a deficiency in the host leading to adverse effects (hematopoiesis and foetological disorders) (14). In addition to these side effects, there is drug resistance (15); therapeutic failures (16, 17), and lack of a human vaccine shows that it is of great importance to identify novel potent candidates that would be well-tolerated and act on both tachyzoites and cysts.

The antiparasitic properties of natural products, such as plant derived substances, have been extensively investigated (18). A review carried out between the period 1981-2019 stated that out of 20 antiparasitic drugs proven as a treatment, there were, nine derived from natural products and three were derived from synthetic pharmacophoric sources of natural products (19). In recent years, several studies have been conducted *in vitro* and or *in vivo* that have shown that plant species could effectively inhibit the growth of *T. gondii* (15, 20, 21).

In the course of continuing research for bioactive compounds from medicinal plants and based on a literature survey on plants traditionally used to treat bacterial and parasitic diseases in Côte d’Ivoire, eight indigenous plant species were selected among those most widely used by traditional healers and for which anti-*toxoplasma* activity had never been evaluated: *Combretum micranthum* G. Don (Combretaceae) leaves, *Elaeis guineensis* Jack. (Arecaceae) leaves, *Erigeron floribundus* Kunth Sch.Bip (Asteraceae) aerial parts, *Oldfieldia africana* Benth. & Hook. F. (Picrodendraceae) stem bark, *Octoknema borealis* Hutch. & Dalziel (Olacaceae) leaves and stem bark, *Omphalocarpum ahia* A. Chev. (Sapotaceae) leaves and stem bark, *Omphalocarpum elatum* Miers stem bark, and whole plant of *Tristemma coronatum* Benth (Melastomataceae).

*C. micranthum*, commonly called Kinkeliba, is an African medicinal plant with several ethnopharmacological activities and used in West Africa as a traditional medication for the treatment of malaria, diabetes, high blood pressure and inflammatory diseases. Its leaves extracts showed antitoxoplamosis (18), anti-diabetic (22), antibacterial (23), antiviral activity (24), antioxidant (25), antityrosinase (26) and protective effect against kidney damage (27). *E. guineensis*, or palm oil tree originates from West and Central Africa, has been reported as a traditional medicine for wound healing, cancer, headaches, rheumatism, and is considered as diuretic. Its leaves extracts showed antioxidant (28−30), anti-plasmodial (31) and promotes vascular relaxation (32). *E. floribundus* is a terrestrial herbaceous plant. In Côte d’Ivoire, this plant is used for the treatment of skin disorders, dyspepsia, abdominal pains, various diseases of microbial origin (33), in AIDS therapy, rheumatism, gout, cystitis, nephritis, dysmenorrhoea, dental pain, headache (34, 35). The aerial parts extracts showed antioxidant, antimicrobial (36), antifungal (37), anthelminthic (38), anti-*Trypanosoma brucei* (39), antimalarial (40), and immunomodulatory effect (41). *O. africana* is a large tree considered in Côte d’Ivoire as a powerful fetish tree with efficacious medicinal virtues. The bark is antiseptic and haemostatic and decoction is used to treat blennorrhoea and act as a pelvic decongestant (35). The stem bark extracts showed antimalarial (42), antibacterial and antifungic activities (43). *O. borealis*, is a large tree. The stem bark is used in traditional medicine for the treatment of lung disorders and to cure coughs whereas the leaves are used as febrifuges. The pulverized bark is rubbed on the skin as a treatment for fever (35, 44). *O. ahia* is an evergreen medium-sized tree. It is used in folklore medicine for the treatment of pain, inflammation, bacterial and parasitic diseases (45, 46). The stem bark extracts showed antibacterial, anti-leishmania and anti-inflammatory activities (47). *O. elatum* is a large tree from West African, used traditionally for the treatment of sterility of males and increases lactation in women. A decoction of the bark, combined with the fruits of peppers (*Capsicum annuum*) and *Solanum anguivi*, is used to treat malaria. A decoction of the young leaves is used for the treatment of cough. Its stem bark extracts showed high *in vitro* anthelmintic (48), anti-inflammatory and antimicrobial activities (49). *T. coronatum* is a semi-prostrate herb or shrub to nearly 1 m high in the ground vegetation of the closed forest, from Guinea to Togo. Leaves are used in folklore medicine as pain-killers, sedatives, for arthritis and rheumatism (35), against typhoid fever, hemorrhoids, infertility, and skin diseases (50). Its leaves extracts showed antibacterial activities (51).

In this study, *n*-heptane (*n*-Hept), ethyl acetate (EtOAc), dichloromethane (DCM) and aqueous methanol 80% (MeOH 80%) extracts were prepared from the ten selected samples and tested *in vitro* at 25 μg/mL to evaluate their activity against *T. gondii*, and cytotoxicity on normal Vero cell line. Then a ^13^C-NMR-based dereplication workflow combined with a bioactivity-guided fractionation process was applied on the most active extract (MeOH 80% extract of the leaves of *E. guineensis*) to identify the compounds being involved in the biological effect against *T. gondii*.

## RESULTS

### Biological screening of crude extracts *on T. gondii*

The *n-*Hept, DCM, EtOAc and 80%MeOH extracts of the ten selected drugs were evaluated for their cytotoxic activity against Vero cells. Among the 40 tested extracts, six were found to be cytotoxic at the concentration of 25 μg/mL, namely the *n*-Hept extract of *T. coronatum,* the DCM extracts of *E. floribundus* and of *O. ahia* leaves, the EtOAc extracts of leaves and bark of *O. borealis* and the MeOH 80% extract of *O. elatum* (Table 1). Therefore, only the non-cytotoxic extracts were then evaluated for their antitoxoplasmic activity *in vitro* at 25 μg/mL. The results (Table 1) of the anti-*Toxoplasma* screening revealed that eight of the 34 non-cytotoxic extracts showed more than 50% parasite growth inhibition at 25µg/mL. These extracts were the DCM and EtOAc extracts of *C. micranthum*, the MeOH 80% extract of *E. guineensis*, the *n*-Hept and DCM of *O. africana*, the EtOAc of the leaves and DCM of the stem bark of *O. ahia*, the DCM extract of *O. elatum* and finally the DCM extract of *T. coronatum* with a respective parasite growth inhibition rate of 79%, 64%, 94%, 67%, 87%, 91%, 96%, 57% and 72%. The nine bioactive extracts are obtained from six different plant species. The hydromethanolic extract of *E. guineensis* leaves was selected for further study in the rest of the study.

**Table 1.**
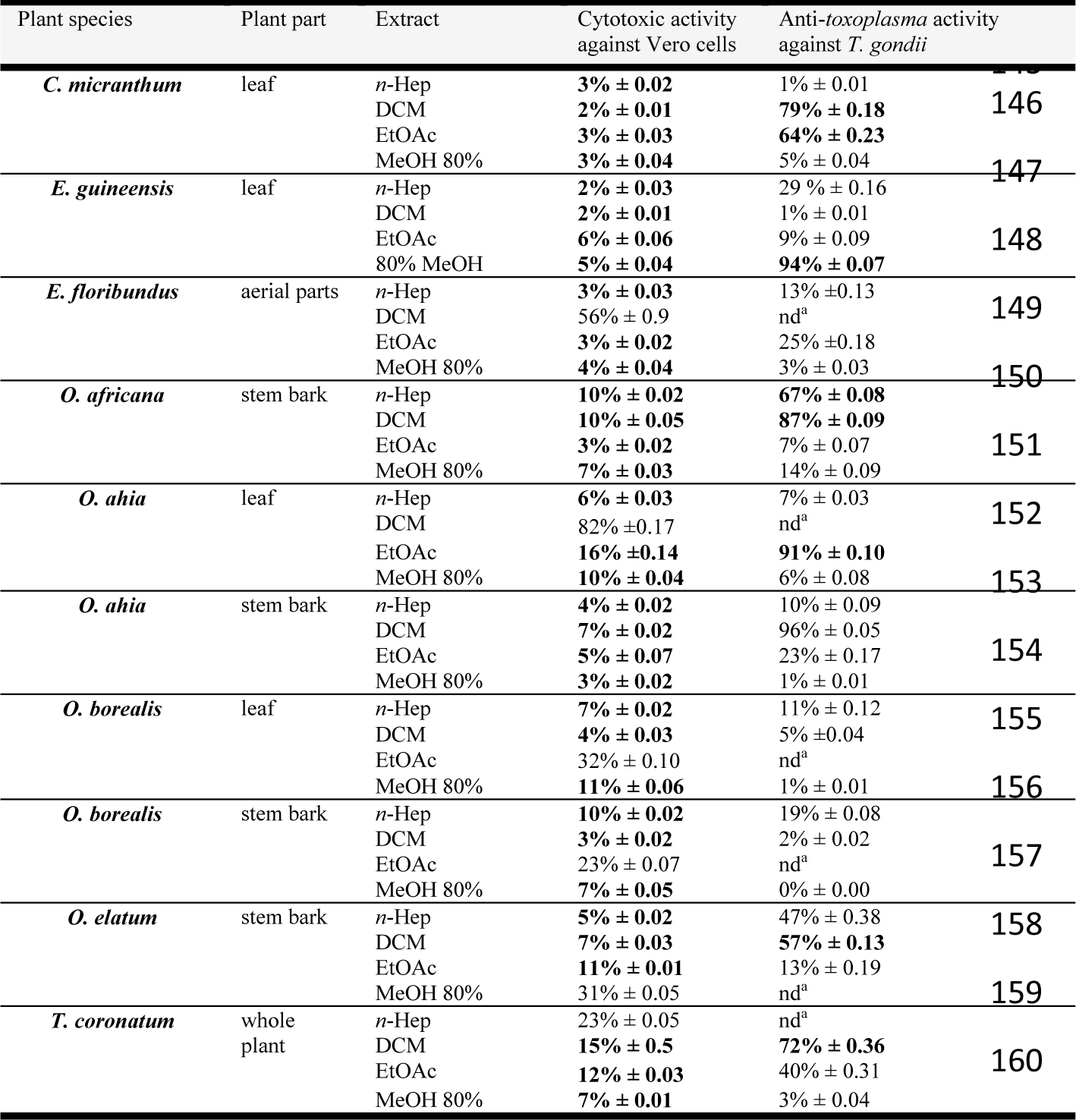
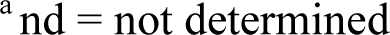
*In vitro* cytotoxicity towards Vero cells for the extracts of the selected plants Vero cells and the *in vitro* anti-*Toxoplasma* activity of extracts exhibiting less than 20% cytotoxicity at 25 µg/mL (N=6).

| Plant species | Plant part | Extract | Cytotoxic activity against Vero cells | Anti-*toxoplasma* activity against *T. gondii* |  |
| --- | --- | --- | --- | --- | --- |
| *C. micranthum* | leaf | *n*-Hep | 3% ± 0.02 | 1% ± 0.01 | 146 |
|  |  | DCM | 2% ± 0.01 | 79% ± 0.18 |  |
|  |  | EtOAc | 3% ± 0.03 | 64% ± 0.23 |  |
|  |  | MeOH 80% | 3% ± 0.04 | 5% ± 0.04 |  |
| *E. guineensis* | leaf | *n*-Hep | 2% ± 0.03 | 29% ± 0.16 | 147 |
|  |  | DCM | 2% ± 0.01 | 1% ± 0.01 |  |
|  |  | EtOAc | 6% ± 0.06 | 9% ± 0.09 |  |
|  |  | 80% MeOH | 5% ± 0.04 | 94% ± 0.07 |  |
| *E. floribundus* | aerial parts | *n*-Hep | 3% ± 0.03 | 13% ± 0.13 | 149 |
|  |  | DCM | 56% ± 0.9 | nd<sup>a</sup> |  |
|  |  | EtOAc | 3% ± 0.02 | 25% ± 0.18 |  |
|  |  | MeOH 80% | 4% ± 0.04 | 3% ± 0.03 |  |
| *O. africana* | stem bark | *n*-Hep | 10% ± 0.02 | 67% ± 0.08 | 150 |
|  |  | DCM | 10% ± 0.05 | 87% ± 0.09 |  |
|  |  | EtOAc | 3% ± 0.02 | 7% ± 0.07 |  |
|  |  | MeOH 80% | 7% ± 0.03 | 14% ± 0.09 |  |
| *O. ahia* | leaf | *n*-Hep | 6% ± 0.03 | 7% ± 0.03 | 152 |
|  |  | DCM | 82% ± 0.17 | nd<sup>a</sup> |  |
|  |  | EtOAc | 16% ± 0.14 | 91% ± 0.10 |  |
|  |  | MeOH 80% | 10% ± 0.04 | 6% ± 0.08 |  |
| *O. ahia* | stem bark | *n*-Hep | 4% ± 0.02 | 10% ± 0.09 | 153 |
|  |  | DCM | 7% ± 0.02 | 96% ± 0.05 |  |
|  |  | EtOAc | 5% ± 0.07 | 23% ± 0.17 |  |
|  |  | MeOH 80% | 3% ± 0.02 | 1% ± 0.01 |  |
| *O. borealis* | leaf | *n*-Hep | 7% ± 0.02 | 11% ± 0.12 | 155 |
|  |  | DCM | 4% ± 0.03 | 5% ± 0.04 |  |
|  |  | EtOAc | 32% ± 0.10 | nd<sup>a</sup> |  |
|  |  | MeOH 80% | 11% ± 0.06 | 1% ± 0.01 |  |
| *O. borealis* | stem bark | *n*-Hep | 10% ± 0.02 | 19% ± 0.08 | 156 |
|  |  | DCM | 3% ± 0.02 | 2% ± 0.02 |  |
|  |  | EtOAc | 23% ± 0.07 | nd<sup>a</sup> |  |
|  |  | MeOH 80% | 7% ± 0.05 | 0% ± 0.00 |  |
| *O. elatum* | stem bark | *n*-Hep | 5% ± 0.02 | 47% ± 0.38 | 158 |
|  |  | DCM | 7% ± 0.03 | 57% ± 0.13 |  |
|  |  | EtOAc | 11% ± 0.01 | 13% ± 0.19 |  |
|  |  | MeOH 80% | 31% ± 0.05 | nd<sup>a</sup> |  |
| *T. coronatum* | whole plant | *n*-Hep | 23% ± 0.05 | nd<sup>a</sup> | 159 |
|  |  | DCM | 15% ± 0.5 | 72% ± 0.36 |  |
|  |  | EtOAc | 12% ± 0.03 | 40% ± 0.31 |  |
|  |  | MeOH 80% | 7% ± 0.01 | 3% ± 0.04 |  |

<sup>a</sup> nd = not determined

### Bioguided screening of MeOH extract of *Elaeis guineensis* on *T. gondii*

Based on the results of the anti-toxoplasmic activity screening, an extraction was carried out on 1.5 kg of the air-dried and powdered *E. guineensis* leaves. The hydromethanolic extract was fractionated on a Diaion HP-20 resin column to obtain HP_1_–HP_5_ fractions. The cytotoxicity of the MeOH 80% extract and obtained fractions was evaluated at 25 μg/mL on Vero cells (Table 2) and showed that the less polar fraction HP_5_ is cytotoxic with a growth inhibition rate of 67%. Therefore, HP_5_ was excluded from the antiparasitic screening. The HP_1_–HP_4_ fractions were evaluated for their anti-*Toxoplasma* activity at the concentration of 25 μg/mL, but none showed anti-*Toxoplasma* activity (Table 2). It seems that the fractionation of the hydromethanolic extract led to the loss of the anti-toxoplasmic activity. A chemosensitivity on the HP_5_ fraction was carried out to determine the concentration at which it is no longer cytotoxic, in order to measure its anti-toxoplasmic activity. The cytotoxicity of HP_5_ was evaluated on Vero cells at 25, 12.5, 6.25, 3.125 and 1.563 μg/mL and the IC_50_ was found to be 16.0 µg/mL (Fig. S1). The concentration which induced less than 12 % of Vero cells growth inhibition was 6.25 µg/mL. Unfortunately, HP_5_ showed less than 1% *T. gondii* growth inhibition at this concentration. The CPC fractionation of the MeOH 80% extract led to eleven fractions which were evaluated for their cytotoxicity on Vero cells at 25 μg/mL showing that the less polar fractions (CPC_1_– CPC_3_) possessed cytotoxic activity (Table 2). The non-cytotoxic fractions were evaluated for their antiparasitic activity, unfortunately revealing that CPC_4_– CPC_11_ fractions possessed less than 50% anti-*Toxoplasma* activity at 25 µg/mL.

**Table 2.** Cytotoxicity and anti-*Toxoplasma* activities of fractions obtained from MeOH 80% extract of *E. guineensis* against *T. gondii* in Vero cells at 25 μg/mL (N=10).

| Fractions from hydromethanolic extract of *E. guineensis* | Molecular family | Cytotoxic activity against Vero cells | Anti-*Toxoplasma gondii* activity |
| --- | --- | --- | --- |
| MeOH 80% extract | - | 5% ± 0.04 | 94% ± 0.07 |
| HP<sub>1A</sub> | Saccharides | 7% ± 0.09 | 2% ± 0.02 |
| HP<sub>1B</sub> | Glycerolipids | 4% ± 0.08 | 5% ± 0.07 |
| HP<sub>2</sub> | Flavonoids | 6% ± 0.07 | 6% ± 0.10 |
| HP<sub>3</sub> | Flavonoids, terpenoids | 2% ± 0.04 | 8% ± 0.12 |
| HP<sub>4</sub> | Flavonoids, terpenoids and sterols | 3% ± 0.02 | 1% ± 0.02 |
| HP<sub>5</sub> | Terpenoids, flavonoids and fatty acids | 67% ± 0.08 | nd<sup>a</sup> |
| CPC<sub>1</sub> | Fatty acids and glycerolipids | 81% ± 0.07 | nd<sup>a</sup> |
| CPC<sub>2</sub> | Terpenoids and sterols | 77% ± 0.04 | nd<sup>a</sup> |
| CPC<sub>3</sub> | Terpenoids and sterols | 47% ± 0.07 | nd<sup>a</sup> |
| CPC<sub>4</sub> | Sesquiterpenoids | 9% ± 0.17 | 22% ± 0.015 |
| CPC<sub>5</sub> | Sesquiterpenoids | 8% ± 0.09 | 8% ± 0.07 |
| CPC<sub>6</sub> | Saccharides | 5% ± 0.04 | 0% ± 0.0 |
| CPC<sub>7</sub> | Saccharides | 4% ± 0.04 | 15% ± 0.15 |
| CPC<sub>8</sub> | Flavonoids | 2% ± 0.04 | 1% ± 0.01 |
| CPC<sub>9</sub> | Sesquiterpenoids and glycosides | 3% ± 0.02 | 1% ± 0.01 |
| CPC<sub>10</sub> | Sesquiterpenoids and glycosides | 5% ± 0.10 | 11% ± 0.13 |
| CPC<sub>11</sub> |  | 1% ± 0.01 | 7% ± 0.07 |

<sup>a</sup> nd = not determined

### Chemical profiling by dereplication ^13^C-NMR of the HP-20 fractions of the MeOH 80% extract of E. guineensis and purification of the HP5 fraction

The result therefore showed that *T. gondii* is sensitive to the combination of HP_5_ with HP_2_, HP_3_ or HP_4_. Considering the above results, chemical profiling of the six fractions HP_1A_–HP_5_ by ^13^C-NMR based dereplication was initiated in order to identify the main chemicals that could potentially be involved in the activity of the MeOH 80% extract (Fig. 1). Nine compounds, including two saccharides (**1**, **2**), three flavonoids (**4–6**), one sesquiterpene glucoside (**7**), three aliphatic acid derivatives (**3**, **8**, **9**) were identified. The correlated chemical shifts of cluster 1 in fraction HP_1A_ were assigned to glucopyranose (**1**) and *α*-galactopyranosyl-(1→3)-*β*-galactopyranoside (**2**) (65). Similarly, cluster 2 in fraction HP_1B_ proposed 1-monopalmitin (**3**) (66).

**FIG 1.**
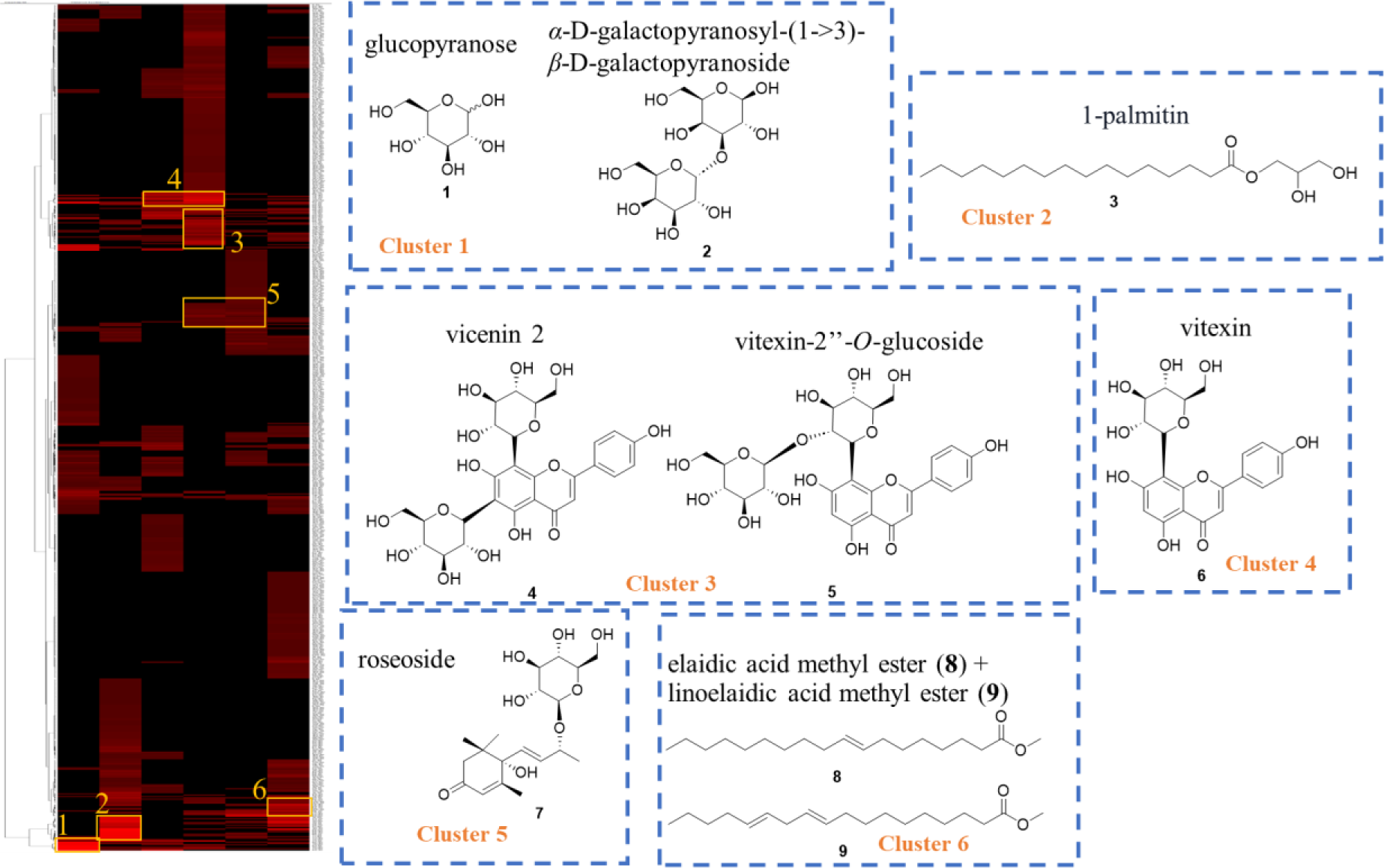
Heatmap of peak intensities of ^13^C NMR (rows) and fractions (columns) from the MeOH 80% extract of *E. guineensis.* This representation allows the visualization of the carbon skeleton of the major compounds.

The cluster 3 in fraction HP_3_ proposed vicenin-2 (**4**) (67) and vitexin-2’’-*O*-*β*-D-glucopyranoside (**5**) (68). The cluster 4 in fraction HP_3_ proposed vitexin (**6**) (69). The cluster 5 in fraction HP_4_ proposed roseoside (**7**) (70) whereas cluster 6 in fraction HP_5_ proposed elaidic acid methyl ester (**8**) and linoelaidic acid methyl ester (**9**) (71) (Fig. 1). The identification of these compounds was confirmed by comparison with literature data and by analysis of HR-ESI-MS spectra of HP_1A_–HP_5_.

In addition, the HP_5_ fraction was purified giving four aliphatic acid derivatives, two cardanol derivatives, one triterpene and one sterol. Their structures were established by using a combination of 1D- and 2D-NMR spectroscopic and mass spectrometry analyses, as well as by comparison with literature values as elaidic acid methyl ester (**8**) and linoelaidic acid methyl ester (**9**), palmitic acid (**10**), palmitic acid methyl ester (**11**) (71), 3-[12(*E*)-pentadecenyl]phenol (**12**), 3-[14(*E*)-heptadecenyl]phenol (**13**) (72), ursolic acid (**14**) (73) and stigmasta-4,24(28)-dien-3-one(**15**) (74) (Fig. 2).

**Fig 2.**
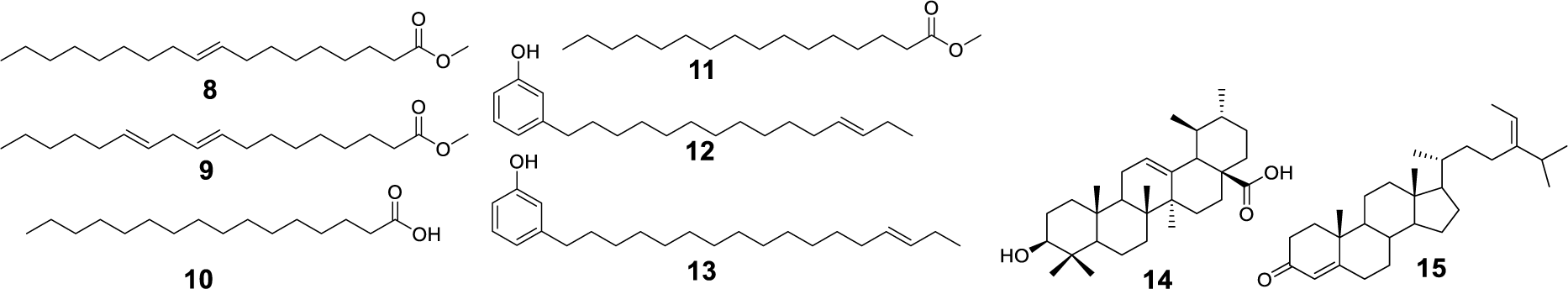
Structure of compounds (**8–15**) purified from HP_5_ fraction of *E. guineensis*.

## DISCUSSIONS

The toxicity induced by these extracts on cells leads to cell death, and therefore would have as an immediate consequence preventing cell invasion by tachyzoites or parasite multiplication. As a result, during the ELISA enzyme-linked immunosorbent assay, very few parasites will be labeled by antibodies, which will lead to a low “false” optical density (OD) result (represents the concentration of parasites present in each well), which does not reflect the true inhibition of the drug on parasite multiplication. Therefore, only the non-cytotoxic extracts were then evaluated for their antitoxoplasmic activity *in vitro* at 25 μg/mL.

The antitoxoplasmic activity of *C. micranthum* has already been described (57) and previous study showed that its aqueous extract had moderate antitoxoplasmic activity with IC_50_ 217 μg/mL (18) and 254 μg/mL (58) against RH tachyzoites of *T. gondii* in the RMC5 cells. However, our work showed that the low-medium polar leaves extracts: DCM and EtOAc of this plant possessed antitoxoplasmic activity at 25 μg/mL on Vero cells with 79% ± 0.18, and 64% ± 0.23 growth inhibition, respectively. The use in traditional medicine of the genus *Omphalocarpum* as an antiparasitic is more highlighted in the literature. This fact has been corroborated by our results which supported the traditional use of the stem bark of *O. ahia* as an antiparasitic (46) based on the strong inhibition of *T. gondii* growth at 25 μg/mL of the DCM extracts from stem bark (96% ± 0.05) and EtOAc from leaves (91% ± 0.10). Whereas DCM extract of *O. elatum* was moderately active against *T. gondii* tachyzoites (57% ± 0.13). In traditional medicine, the stem bark of *O. africana* are boiled with palm oil to make an ointment to treat infestations against lice (a parasitic disease). Our work showed that its *n*-Hep and DCM stem bark extracts were active against *T. gondii* at 25 μg/mL with 67% ± 0.08, and 87% ± 0.09 growth inhibition, respectively. *Tristemma coronatum* is used in traditional medicine as antibacterial (51). Our work showed that it has also an antiparasitic activity. The traditional antiparasitic use of *E. guineensis* leaves in Côte d’Ivoire that had been observed for the 70% hydroethanolic extract of leaves against *P. falciparum (*another *Apicomplexa parasite)* with IC_50_ 1.195 μg/mL (31), was supported by the strong inhibition (94 % ± 0.07) of *T. gondii* growth at 25 μg/mL by the hydromethanolic 80% extract. We therefore selected this extract for further study to identify the compounds being involved in the biological effect against *T. gondii*.

It is important to point out that, despite the historical effectiveness of bioguided fractionation of plant extracts, loss of activity and failure to isolate active compounds during this process can occur (59, 60). The main reasons of these pitfalls may be the degradation of compounds during the purification process, low concentration of bioactive compounds making their isolation difficult, and/or potential synergistic effects (60). This can be avoided by “synergy-directed fractionation” (61) which combines chromatographic separation and synergy testing in combination with a known active constituent in the original extract. Several studies have shown that the overall activity of plant extracts can result from mixtures of compounds with synergistic, additive, or antagonistic activity (62).

While there are multiple possible explanations for this failure, it is true that in some cases loss of activity occurs because multiple constituents are required to observe the biological effect. The first explanation could be the irreversible adsorption of compounds on the column packing (63). To test this purpose, the MeOH 80% extract was fractionated by centrifugal partition chromatography (CPC), a solid support free liquid-liquid separation technique involving the distribution and transfer of solutes between at least two immiscible liquid phases according to their partition coefficients. This method avoids irreversible adsorption of analytes due to the absence of solid support. Unfortunately, its fractionation by CPC had not yet made it possible to find the anti-*T. gondii* activity in its individualized fractions. Thus, the loss of activity of the MeOH 80% extract is not due to irreversible adsorption of compounds to the column. This suggests that the anti-*toxoplasma* activity of the MeOH 80% extract could result from mixtures of compounds with synergistic, additive, or antagonistic activity.

The second hypothesis was that there was a synergistic interaction between the metabolites of the MeOH 80% extract. It is known that when two or more drugs are given in combination, the effect can be superadditive (synergistic); that is, they can demonstrate action beyond that is expected of their individual power and effectiveness. Unlike synergy, some drug combinations may show sub-additivity (antagonism) or simple additivity (64). Thus, the idea that HP_5_, the only cytotoxic fraction at 25 μg/mL, could be responsible of the anti-*toxoplasma* activity of the MeOH 80% extract of *E. guineensis* at lower concentrations when mixed with another fraction was verified. So, five combinations were reconstituted HP_1A_+HP_5_, HP_1B_+HP_5_, HP_2_+HP_5_, HP_3_+HP_5_ and HP_4_+HP_5_, respecting their ratios in the MeOH 80% extract (Table 3). These combinations were evaluated for their cytotoxicity at the concentration of 25 µg/mL on Vero cell line (Table 3) and, surprisingly, none of these combinations showed more than 20% cytotoxic activity at the concentration tested. The combination tested HP_1B_ + HP_5_ (25 µg/mL), in which the HP_5_ concentration was 13.88 µg/mL showed a toxicity of 19%± 0.01 while HP_5_ tested alone showed toxicity of 40% at this concentration (Fig. S1). The other combinations tested HP_2_+HP_5_, HP_3_ + HP_5_ and HP_4_ + HP_5_ also showed that these combinations were less cytotoxic than HP_5_ alone at the same concentration for HP_5_ (Table 3). These results suggested a sub-additive (antagonist) effect concerning the cytotoxic activity on Vero cell line.

**Table 3.** Cytotoxicity and anti-*Toxoplasma* activities of some fractions and combinations of MeOH 80% extract of *E. guineensis.*

| Fractions | Concentration in $\mu\text{g/mL}$ | Cytotoxicity against Vero cells | Anti-*toxoplasma* activity at 25 $\mu\text{g/mL}$ | Anti-*toxoplasma* activity at 100 $\mu\text{g/mL}$ |
| --- | --- | --- | --- | --- |
| HP<sub>1B</sub> | 11.1 | nd<sup>a</sup> | nd<sup>a</sup> | nd<sup>a</sup> |
| HP<sub>5</sub> | 13.9 | 40% ± 0.01 | nd<sup>a</sup> | nd<sup>a</sup> |
| HP<sub>1B</sub>+HP<sub>5</sub> | 25 | **19% ± 0.01** | **64% ± 0.05** | **92% ± 0.02** |
| HP<sub>2</sub> | 14.2 | nd<sup>a</sup> | nd<sup>a</sup> | nd<sup>a</sup> |
| HP<sub>5</sub> | 10.8 | 30% ± 0.01 | nd<sup>a</sup> | nd<sup>a</sup> |
| HP<sub>2</sub>+HP<sub>5</sub> | 25 | **7% ± 0.05** | **67% ± 0.03** | **85% ± 0.03** |
| HP<sub>4</sub> | 16.9 | nd<sup>a</sup> | nd<sup>a</sup> | nd<sup>a</sup> |
| HP<sub>5</sub> | 8.1 | 20% ± 0.01 | nd<sup>a</sup> | nd<sup>a</sup> |
| HP<sub>4</sub>+HP<sub>5</sub> | 25 | **10% ± 0.07** | **74% ± 0.02** | **92% ± 0.03** |
| HP<sub>3</sub> | 19 | nd<sup>a</sup> | nd<sup>a</sup> | nd<sup>a</sup> |
| HP<sub>5</sub> | 6 | **15% ± 0.01** | 1% ± 0.00 | nd<sup>a</sup> |
| HP<sub>3</sub>+HP<sub>5</sub> | 25 | **2% ± 0.01** | **50% ± 0.08** | **83% ± 0.05** |
| HP<sub>1A</sub> | 23.5 | nd<sup>a</sup> | nd<sup>a</sup> | nd<sup>a</sup> |
| HP<sub>5</sub> | 1.5 | 0 ± 0.00 | nd<sup>a</sup> | nd<sup>a</sup> |
| HP<sub>1A</sub>+HP<sub>5</sub> | 25 | **3% ± 0.01** | **0 ± 0.00** | **0 ± 0.00** |

<sup>a</sup> nd = not determined

These five combinations were evaluated for their anti-*Toxoplasma* activity at the concentration 25 µg/mL. The HP_3_+HP_5_ combination in which the concentration of HP_5_ was 5.95 µg/mL showed an anti-*Toxoplasma* effect of 50%± 0.08, resulting in synergistic activity since the anti-*Toxoplasma* activity of HP_5_ alone at the same concentration was 1%. The most active combination was HP_4_ + HP_5_ with 74% ± 0.02 antiparasitic activity (Table 3). It is believed that a synergistic effect between the identified (**1–9**) and purified (**8−15**) metabolites of these fractions would be responsible for the anti-Toxoplasma activity. However, further studies should be carried out in order to confirm and explain this synergistic or antagonistic effect between the identified compounds. All CPC fractions (CPC_1_−CPC_11_) were analyzed by ^1^H and ^13^C NMR spectroscopy in order to identify the main class of compounds (Table 2) and compared to HP_1A_–HP_5_ fractions. No additional compounds were identified. The activity observed in this plant could be attributed to a synergistic effect between the phytochemical constituents, including aliphatic acids, terpenoids, sterols and flavonoids which could have antiparasitic activities. However, further studies should be carried out to confirm and explain this synergistic or antagonistic effect between the identified compounds.

Fatty acids are a broad group of amphipathic molecules with a terminal carboxyl group that, based on the number of double bonds, are classified into saturated and unsaturated fatty acids. These molecules play a crucial role in the structure of cell membranes, energy production, signaling, and regulation of different biochemical pathways (75). They are commonly found at cell membranes, and act as second messengers that can modify intracellular and extracellular signaling pathways, affecting gene expression and physiological and metabolic responses in different tissues (76−78). Long-chain fatty acids contained in HP_5_ were palmitic acid, palmitic acid methyl ester, elaidic acid methyl ester and linoelaidic acid methyl ester. Three of them belong to fatty acid methyl esters (FAMEs) which have been reported to exhibit antibacterial properties. Some authors have even described that FAMEs could be the next generation of antibiotic agents acting through membrane disruption and reactive oxygen species production (79). Most likely, fatty acids from the active fraction HP_5_ affected the lipid bilayer integrity or even the inner membrane complex of *T. gondii*. Both structures are vital for invasion and replication of the parasite (80). Similar compounds, such as hexadecanoic acid methyl ester, have been shown to affect the integrity of bacterial cell membranes (81). Similarly, the methyl esters of 9-octadecenoic, octadecanoic and hexadecanoic acids were identified are the main constituents of diverse medicinal plants, such as *Gracilaria* spp., *Leucaena leucocephala*, and *Salix babylonica* (15, 82, 83). In addition to affecting cell membranes, inhibition of bacterial fatty acid synthesis is suggested to be linked to these molecules (84, 85). *T. gondii* acyl-CoA diacylglycerol acyltransferase (TgDGAT) is an integral membrane protein localized in the cortical and perinuclear endoplasmic reticulum of the parasite and is responsible for triacylglycerol synthesis (15). Some studies have reported that oleate, palmitoleate, and linoleate impair parasite replication by interfering with TgDGAT function (86). Therefore, *T. gondii* growth can be limited by controlling the parasite lipidomics, as previously reported (15, 87). Based on these reports, the fatty acids in HP_5_ could disrupt the parasite’s membranes or inhibit enzymes required for *T. gondii* fatty acid synthesis.

Terpenes are characterized by their isoprene unit number and have been extensively reported to exhibit antibiotic, anticancer, antioxidant, antiviral, antihyperglycemic, and antiparasitic properties (88−90). Triterpenoids commonly observed as natural products, ursolic acid and stigmasta-4,24(28)-dien-3-one, were detected in HP_5_. The growth inhibitory activity of these metabolites against *T. gondii* has already been reported with IC_50_ of 5.9 µM for ursolic acid (91−94) and 17.3 µM for *β*-sitostenone, the analogue of stigmasta-4,24(28)-dien-3-one (21). Illustrating their promise as a potential candidate for anti-*Toxoplasma gondii* drug development, lupane-type triterpenes extracted from the bark of *Alnus glutinosa* showed anti-*Toxoplasma* activity *in vitro*. Betulone was the most active triterpene (IC_50_ 2.7 µM) (20). Triterpenoids isolated from the barks of *Quercus crispula*, 29-norlupane-3,20-dione, oleanolic acid acetate, and ursolic acid acetate, showed anti-*Toxoplasma* activity with an IC_50_ between 6.8 and 24.4 Μm (93). A pentacyclic triterpenoid from *Olea europaea*, maslinic acid, was evaluated on *T. gondii* tachyzoites and showed an IC_50_ of 58.2 μM. It seems that this terpenoid disrupts the ultrastructure and motility of the parasite due to inhibition of protease activity (95). Their ability to affect lipid membranes has also been reported (96). Some pentacyclic triterpenes have been shown to inhibit the expression of COX-2 (97), which reduces *T. gondii* infection and upregulates the proinflammatory immune response in mice (98). COX-2 is a crucial factor in *T. gondii* propagation in human trophoblasts; therefore, its inhibition can induce a proinflammatory response capable of controlling parasite proliferation (10). Ursolic acid identified in HP_5_ is also pentacyclic; therefore, it could affect the host Th1 immune response.

*T. gondii* inhibits production of TNF-*α* and IL-12. TNF-*α* is a cytokine of inflammatory and immune response and together with IL-6 can enhance proliferation and differentiation of B lymphocytes [99]. TNF-*α* activates eosinophil cytotoxicity toward protozoa and induces secretion of acute phase proteins via IL-6 production. TNF-*α* and IFN-*γ* have antiproliferative properties. In toxoplasmosis, TNF-*α* appears to be indispensable for macrophage activation and inhibition of parasite replication. Several studies reported the anti-inflammatory properties of *Elaeis guineensis* leaf extracts (100−102). *E. guineensis* extracts contained flavonoids which have antioxidant, apoptosis-inducing, and anti-inflammatory properties (103) and also decreases oxidative stress (103, 104). Flavonoid’s mechanism is based on the activation of nuclear factor Nrf2 (106, 107), which activates other cellular defense mechanisms by Phase II initiation of detoxifying and antioxidant.

In conclusion, our experiments revealed the *in vitro* anti-toxoplasma activities of the DCM and EtOAc extracts of the stem bark of *C. micranthum,* of the MeOH 80% extract of the leaves of *E. guineensis, n*-Hep and DCM extracts from the stem bark of *O. africana,* EtOAc extract from the leaves and DCM extract from the stem bark of *O. ahia*, from the DCM extract of the stem bark of *O. elatum* and the DCM extract of the aerial parts of *T. coronatum*. Further research will need to be investigated to identify their active components and to search new compounds to treat toxoplasmosis. The fractionation of one of the most active extracts, the MeOH 80% extract from the leaves of *E. guineensis*, did not allow recovery of antitoxoplasmic activity in its individualized fractions. After chemical profiling using a ^13^C NMR-based dereplication workflow, the nine major molecular structures were directly identified, highlighting the presence of saccharides (**1-2**), megastigmanes (**7**), flavonoids (**4-6**), terpenes and aliphatic acids (**3, 8, 9**). The purification of one of these fractions led to the identification of palmitic acid (**10**), palmitic acid methyl ester (**11**), 3-[12(*E*)-pentadecenyl]phenol (**12**), 3-[14(*E*)-heptadecenyl]phenol (**13**), ursolic acid (**14**) and tigmasta-4,24(28)-dien-3-one (**15**). Among the compounds elucidated by dereplication or after purification of HP_5_, only vicenin-2 (**4**) and vitexin (**6**) were previously identified in *E. guineensis* by UHPLC-UV/PDA (108, 109).

A synergistic effect between these metabolites is thought to be responsible for its antitoxoplasmic activity. We concluded that MeOH 80% extract of *E. guineensis* is active against *T. gondii* RH and is not cytotoxic to Vero cells at concentrations at 25 µg/mL that killed up to 94% of the parasites. Its efficacy could be related to the mechanisms of action of fatty acids and terpenes; nevertheless, as different molecules are found in this extract, a synergistic or antagonistic effect cannot be discarded. It is therefore crucial to evaluate the components of the MeOH 80% extract as well as their mode of action, both in the parasite and host cells, to validate the vital targets (preferably those not supplied by the host) leading to the identification of a main molecule which will probably allow the design of analogous triterpenoids or FAMEs with anti-*Toxoplasma* activity. Finally, these results demonstrated that *E. guineensis* metabolites can be considered as possible candidates to obtain lead compounds against *T. gondii* and that natural compounds can be a good source for the development of new antiparasitic drugs.

## MATERIALS AND METHODS

### Chemicals, reagents, parasites and cell culture

All solvents for extraction, fractionation and solubilized were purchased from Carlo Erba Reactifs SDS (Val de Reuil, France). The RH strain (genotype I) of *T. gondii* was provided by the Toxoplasma Biological Resource Center (BRC Toxoplasma, France). Tachyzoites of *T. gondii* the RH strain were cultured on Vero cells monolayers (ATCC, CCL81) at 37 °C, 5% CO_2_ in a humidified incubator. Cells and parasites were cultured in Iscove’s Modified Dulbecco’s Medium (IMDM) (Invitrogen, France) supplemented with 5% fetal bovine serum (SVF) for cells cultures or 2% (v/v) SVF for parasites cultures (Biowest, France) and supplemented with 1% antibiotics (penicillin-streptomycin) (GIBCO).

### Plant material

The selected plants were all harvested in February 2021 in their natural habitats in Côte d’Ivoire. Botanical determination was performed by Mr Téré H.G. and authentication was made by Pr. Tiebre S.-M. from the National Floristic Centre (NFC) of university Félix HOUPHOUET-BOIGNY Cocody Abidjan (Côte d’Ivoire), in which, voucher specimens were deposited (Table 4 and Fig. S10).

**Table 4.**
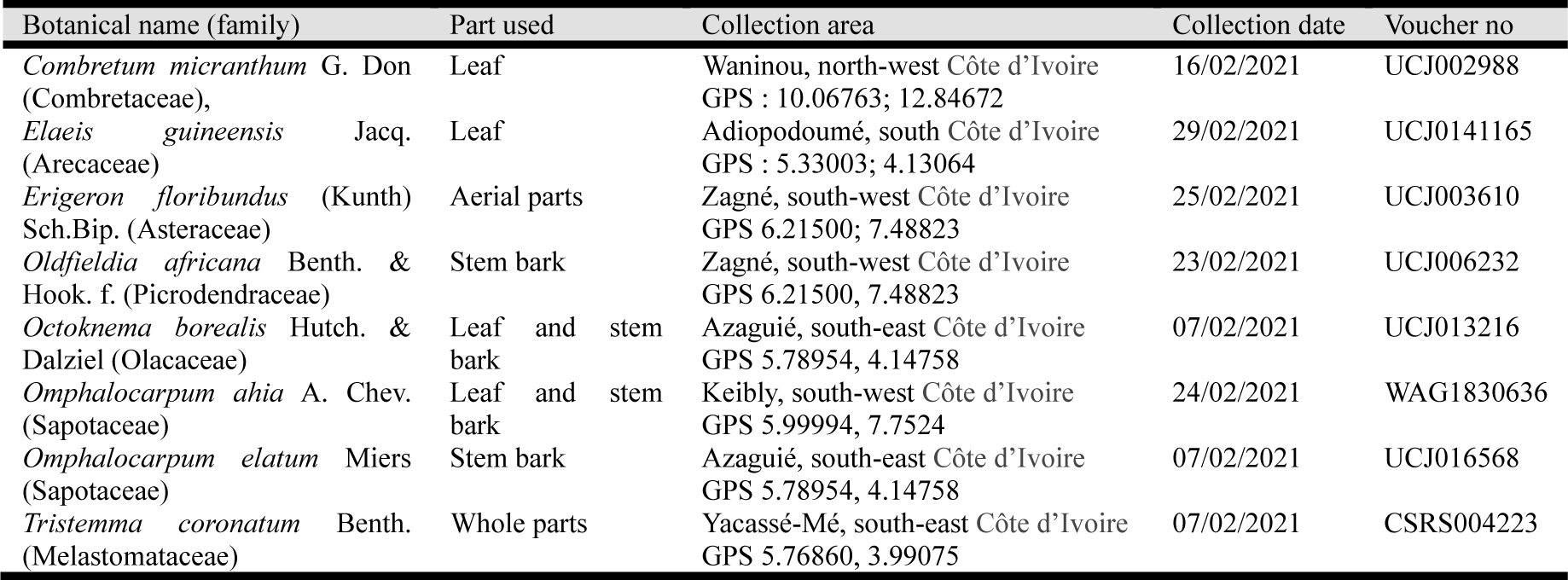
Plant species analyzed for their anti-*Toxoplasma* activity.

### Instrumentation

1D and 2D-NMR spectra were recorded in CD_3_OD on Bruker Avance DRX III 600 instrument using standard Bruker microprograms (Karlsruhe, Germany) equipped with a TCI cryoprobe. The ^13^C NMR spectra were acquired at 150.91 MHz using a standard zgpg pulse sequence with an acquisition time of 0.9 s, a relaxation delay of 3 s. HR-ESI-MS were obtained from Micromass Q-TOF micro-instrument (Manchester, UK). Centrifugal partition chromatography (CPC) experiments were carried out using a lab-scale FCPE300® column of 303 mL capacity (Rousselet Robatel Kromaton, Annonay, France) containing 7 circular partition disks and engraved with a total of 231 partition twin-cells (≈ 1 mL per twin cell). The liquid phases were pumped by a Knauer Preparative 1800 V7115 pump (Berlin, Germany). The column was coupled on-line with a UVD 170 S detector set at 210, 254, 280 and 366 nm (Dionex, Sunnivale, CA, USA). Fractions of 20 mL were collected by a Pharmacia Superfrac collector (Uppsala, Sweden). The column rotation speed was set at 1200 rpm and the flow rate at 20 mL/ min. Flash chromatography was performed on a Grace Reveleris system equipped with dual UV and ELSD detection using Grace cartridges (silica gel or RP-C_18_), and the monitored wavelengths were at 205, 254, and 366 nm. Semi-preparative HPLC was performed on an apparatus equipped with an ASI-100 Dionex autosampler, an Ultimate 3000 pump ThermoFisher Scientific, a diode array detector UVD 340S and a Chromeleon software (Dionex, ThermoFisher Scientific, France). RP-C_18_ column (Phenomenex 250 ×15 mm, Luna 5 µ, Interchim, France) was used for the semi-preparative HPLC with a binary gradient eluent (H_2_O pH 2.4 with TFA; CH_3_CN) and a flow rate of 4.7 mL/min; the chromatogram was monitored at 205, 254, 300, and 360 nm. Analytical TLC was performed using silica gel plates (Merck Kieselgel 60 F254) and RP-C_18_ plates (Kieselgel 60 F_254_) and visualized at 254 and 366 nm and by spraying the dried plates with vanillin and 50 % of H_2_SO_4_, followed by heating.

### Preparation of the 40 crude extracts

Air-dried plant material of each species (10 g) was ground (0.2 mm sieve). The plant material was extracted successively with *n*-hept, DCM, EtOAc and MeOH 80%, at room temperature for 2 × 24 h, each 100 mL. The filtrates were taken to dryness under vacuum at 40°C and the residues were stored at room temperature until testing.

### Extraction and fractionation of *E. guineensis* leaves

Air-dried and powdered leaves of *E. guineensis* (1.5 kg) were extracted, at room temperature, for 2 × 24 h, with *n*-hept, DCM, EtOAc and MeOH 80%, successively, each 15 L, to give after solvent removal under vacuum the corresponding extracts *n*-hept (25.9 g), DCM (13.6 g), EtOAc (7.1 g) and MeOH 80% (44.6 g). A part of the MeOH 80% extract (30 g) was fractionated by Diaion HP20 resin chromatography (5 × 45.5 cm), eluted successively with H_2_O, 25%, 50%, 75% of MeOH in H_2_O and finally with MeOH; each 3 L, to obtain fractions: HP_1_ (20.5 g), HP_2_ (1.5 g), HP_3_ (3.8 g), HP_4_ (2.5 g) and HP_5_ (1.2 g), respectively. Fraction HP_I_ was solubilized in 200 mL of H_2_O and partitioned with *n*-butanol (3 ×250 mL) to obtain the HP_1A_ (H_2_O soluble fraction, 19.6 g) and HP_1B_ (*n*-BuOH soluble fraction, 0.9 g).

### Chemical profiling of the six fractions by ^13^C-NMR based dereplication

Briefly, fractions HP_1_– HP_5_ were chemically profiled based on ^13^C NMR data as described in a previous work (20, 52). All ^13^C NMR spectra of the fraction series were processed and submitted to Hierarchical Clustering Analysis (HCA) for the recognition of ^13^C NMR metabolite fingerprints. In this way, similarity measurements between ^13^C NMR signals belonging to individual structures within the fraction series were visualized as “chemical shift clusters” on a HCA correlation heat map given in Fig. 1. As a result, 9 major chemical shift clusters colored in red were revealed on the heat map, corresponding to the major metabolites of the MeOH 80% extract (Fig. 1). With the help of acd_lotus/nmrshiftdb2 (53) and NP-MRD databases containing chemical shift values of natural metabolites, the corresponding chemical structures were identified.

### Purification of HP_5_ and identification of its major components

Fraction HP_5_ (1.1 g) was separated into 37 subfractions named HP_5-1_−HP_5-37_ using silica gel flash chromatography eluted with *n*-hept-acetone-CHCl_3_-MeOH (from 95:5:0:0 to 0:0:1:1). The combined subfractions HP_5-1_ and HP_5-2_ (21 mg) were purified by semi-prep HPLC, eluted with a gradient from 80% to 100% of CH_3_CN in water as a mobile phase, in 10 min, then in isocratic elution with CH_3_CN for 10 min to afford compounds **14** (*R*_t_ 12.9 min, 1 mg), **8** (*R*_t_ 15.1 min, 1 mg) and **9** (*R*_t_ 18.6 min, 1.3 mg). Subfraction HP_5-3_ (20 mg) was purified by semi-prep HPLC, eluted with a gradient from 80% to 100% of CH_3_CN in water as a mobile phase, in 5 min, then in isocratic elution with CH_3_CN for 20 min to afford compounds **15** (*R*_t_ 16.5 min, 2.1 mg), **13** (*R*_t_ 17.2 min, 1 mg) and **12** (*R*_t_ 23.7 min, 1 mg). Subfraction HP_5-4_ (13 mg) was purified by semi-prep HPLC, eluted with a gradient from 80% to 88% of CH_3_CN in water as a mobile phase, in 10 min to afford compounds **10** (*R*_t_ 7.4 min, 1 mg) and **11** (*R*_t_ 8.9 min, 1 mg).

### Centrifugal partition chromatography (CPC) fractionation of MeOH 80% extract

A three-phase solvent system composed of *n*-Hept/MtBE/CH_3_CN/water in the ratio 1/1/1/1 (v/v/v/v) (500 mL, of each) was used. The solvent mixture was thoroughly equilibrated in a separatory funnel. After separation of the *n*-heptane-rich upper phase (UP_0_), one equivalent volume of MTBE (500 mL) was added to the mixture of middle (MP_0_) and lower (LP_0_) phases in order to slightly reduce the polarity of the remaining two-phase solvent system. After decantation, the final middle and lower phases (MP and LP) were separated. The column was filled with the LP at the minimal rotation speed of 200 rpm. The rotation speed was then increased to 1400 rpm. The sample solution (1.5 g), dissolved in a mixture of LP:MP:UP_0_ (45:10:5 v/v/v), was loaded into the column by progressively pumping the less polar UP_0_ in the ascending mode. The flow rate of the mobile phase was then maintained at 10 mL/min until the end of the experiment. The UP_0_ was pumped for 30 min after the release of the dead volume to ensure the elution of all hydrophobic compounds. The moderately polar MP was then pumped for 30 min to elute compounds with a medium hydrophobicity. Finally, in order to recover the most hydrophilic compounds retained at the head of the column, the role of the two liquid phases was switched by pumping the aqueous LP as the mobile phase in the descending mode at 10 mL/min. Fractions of 20 mL were collected. The separation was monitored by UV at 210, 254, 280 and 366 nm. All fractions were analyzed by TLC and HPLC and then pooled, giving 11 fractions (CPC_1_–CPC_11_).

### Solubilization of extracts and fractions

All extracts and fractions were solubilized in dimethyl sulfoxide (DMSO). The final DMSO concentration was less than 0.5% in the final mixture (DMSO/culture medium, v/v). DMSO is commonly used for *in vitro* and *in vivo* experiments and is an aprotic amphiphilic molecule containing a highly polar domain and two non-polar methyl groups (54). This characteristic allows it to be soluble in polar or non-polar solvents. This also makes it capable of solubilizing many organic compounds polar and non-polar (55, 56).

### Cytotoxic activity

The *in vitro* cytotoxicity of all 40 crude extracts, HP_1_−HP_5_ and CPC_1_–CPC_11_ fractions was assessed on Vero cell cultures using the UptiBlue viable cell count assay (Interchim, France). The crude extracts were prepared at 250 μg/mL in dimethyl sulfoxide (DMSO)/culture medium, v/v, and kept at −30°C, for a final sample tested at 25 μg/mL. A suspension of 20 000 Vero cells were cultured in 96-well plate with IMDM supplemented with 5% SVF (v/v). After 4 hours of incubation, crude extracts or fractions to test were deposited in wells at 25 µg/mL. After 72 hours of incubation at 37°C and 5% CO_2_, wells were washed with phosphate buffer saline and fresh IMDM culture medium supplemented with 5% SVF and 10% UptiBlue was added. After an additional 3 h of incubation, the plate was measured with spectrophotometre with two wavelength, 570 nm corrected at 600 nm (FLUOstar Omega microplate reader, BMG Labtech, France). A cytotoxicity threshold was arbitrarily defined at cell viability of 80%.

### Anti-*Toxoplasma* activity

All the experiments were performed at least in triplicate. The crude extracts were prepared at 250 μg/mL in dimethyl sulfoxide (DMSO)/culture medium, v/v, and kept at −30°C, for a final sample tested at 25 μg/mL. 20,000 Vero cells were filled in 96-well plates with IMDM supplemented with 2% SVF (v/v), then incubated for 3 hours at 37°C and 5% CO_2_. Then, 50,000 parasites of the RH strain contained in 50 μL of culture medium were deposited, followed by 3 h of incubation in the same conditions. This time allowed the parasites to sediment and invade the Vero cells. Host cells were infected in a parasite-to-cell ratio of 5:2. Wells were then washed with PBS and 100 μL and 25 μL of sample solution was deposited per well in duplicate to obtain a final concentration of 25 μg/mL or 100µg/mL. Dilutions were performed with culture medium. Finally, the plates were incubated for 72 h at 37 °C and 5% CO_2_ before being fixed with cold methanol for 5 min at −20 °C and then parasite multiplications were revealed by enzyme immunoassay.

Of the 96 wells, eight wells were reserved as a control of the Vero alone representing 100% cell viability, then six reserved wells were inoculated with *T. gondii* as a parasite growth control under untreated conditions, and finally, two final wells were inoculated under the same conditions as described above with pyrimethamine at 1 μg/mL as a control.

The IC_50_ assays were processed in the same conditions as previously described (21). Cells were then rehydrated with 100 μL of PBS for 10 minutes at room temperature. The plate was treated with 100 μL of a 3% H_2_O_2_ solution for 10 minutes at 37 °C to destroy the endogenous peroxidases of the Vero cells and then was washed with PBS and filled with 60 μL of primary polyclonal anti-*T. gondii* antibody (Bio-rad Rabbit Anti-T. *gondii* 9070-0556) diluted to 1/1000^th^ in a conjugate buffer. After 1 hour of incubation at 37 °C, wells were washed four times with wash buffer. Subsequently, 60 μL of secondary goat antibody anti-rabbit IgG coupled with horseradish peroxidase (Bio-rad Goat Anti-Rabbit IgG (H+L) HRP conjugate 170-6515) diluted to 1/2000^th^ in the conjugate buffer were added, and incubated for 1 hour at 37 °C. The wells were washed four more times and 200 μL of OPD (*o-phenylenediamine dihydrochloride*) substrate (P8412 Sigma-Aldrich, France) was added in each well. After 15 min at room temperature protected from light, the enzymatic reaction was stopped by the addition of 50 μL of hydrochloric acid (HCl) 3 M. To finish, 200 μL of each well were transferred to a new well and a spectrophotometer reading was performed at 492 nm, corrected at 630 nm.

Each experiment was performed with three controls: control with host cells alone, control with host cells and parasites without the drug for parasite multiplication, and control with host cells with parasites and pyrimethamine at 1 μM. The intensity of the staining is proportional to the amount of parasites present in the well.

The anti-*Toxoplasma* activity of samples was determined using the formula: 100 – [Δ_Ods_/ Δ_Odv_) × 100]; Δ_Odv_ = OD at 492 nm – OD at 630 nm for Vero cells invaded by tachyzoites without sample; Δ_Ods_ = OD at 492 nm – OD at 630 nm for Vero cells invaded by tachyzoites and treated with sample.

### Statistical analysis

Statistical significance in plaque assay, proliferation, and fungal growth inhibition assay was evaluated by two-tailed unpaired tests using an Excel spreadsheet. Statistical data are expressed as the mean value ± standard error.

## ACKNOWLEDGEMENTS

The authors thank the ICMR-UMR-7312 CNRS and the UR 7510 ESCAPE of the University of Reims Champagne-Ardenne (France), the National Referencer centre on toxoplasmosis in the CHU of Reims, the UFR Sciences of Structures of Matter and Technology of the Université of Félix-Houphouët-Boigny (Côte d’Ivoire), the Ministry of Higher Education and Scientific Research of Ivory Coast for the financial support.

## AUTHOR CONTRIBUTIONS

**Nangouban Ouattara**: Data curation, Investigation, Methodology, Writing – original draft, Software. **Abdulmagid Alabdul Magid**: Methodology, Supervision, Writing – original draft, Writing – review & editing. **Sandie Escotte-Binet**: Data curation, Investigation, Validation, Supervision. **Philomène Akoua Yao-Kouassi:** Formal analysis, Funding acquisition, Resources, Supervision. **Isabelle Villena^2^**: Funding acquisition, Project administration, Resources, Supervision. **Laurence Voutquenne-Nazabadioko**^1^ Funding acquisition, Project administration, Resources, Supervision, Writing – original draft, Writing – review & editing

## APPENDIX A. SUPPLEMENTARY DATA

The following items are available online at SUPPLEMENTAL FILE 1

---

**FIG S1.**
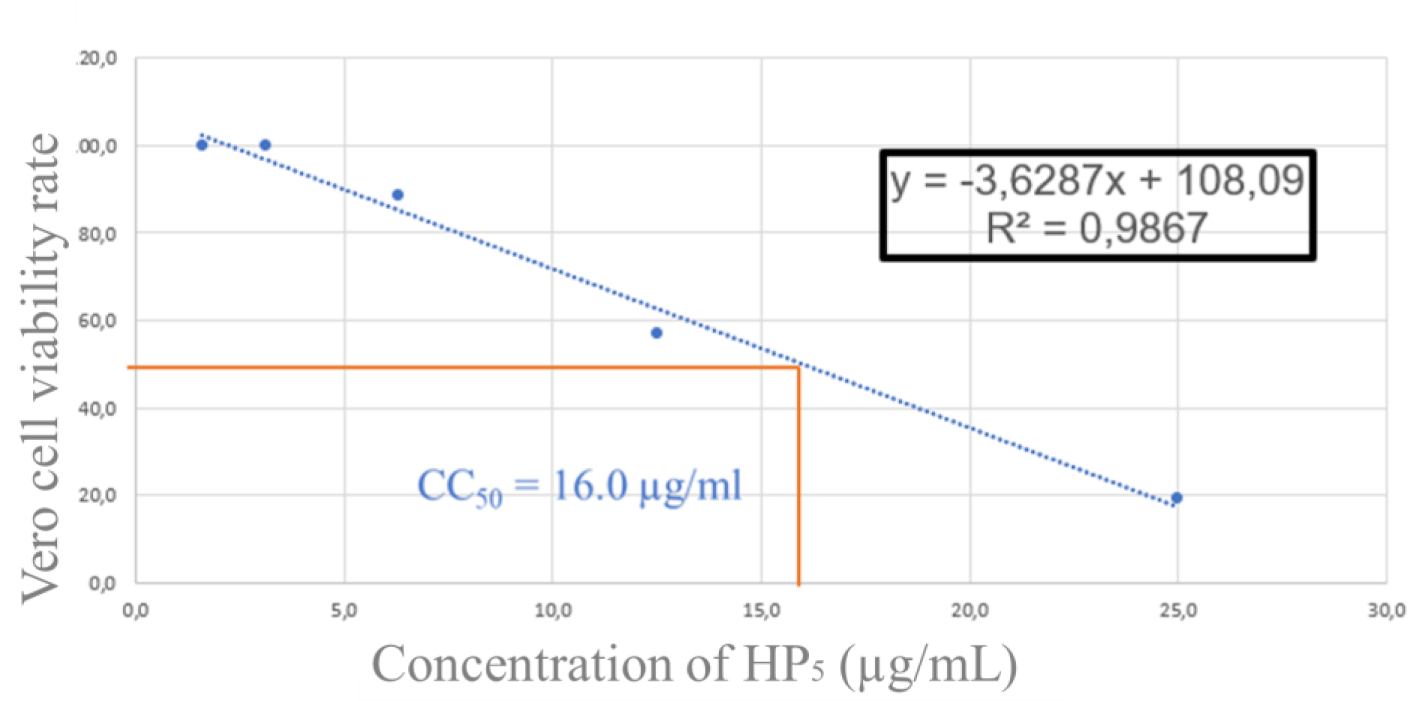
Cytotoxicity of HP_5_

**FIG S2.**
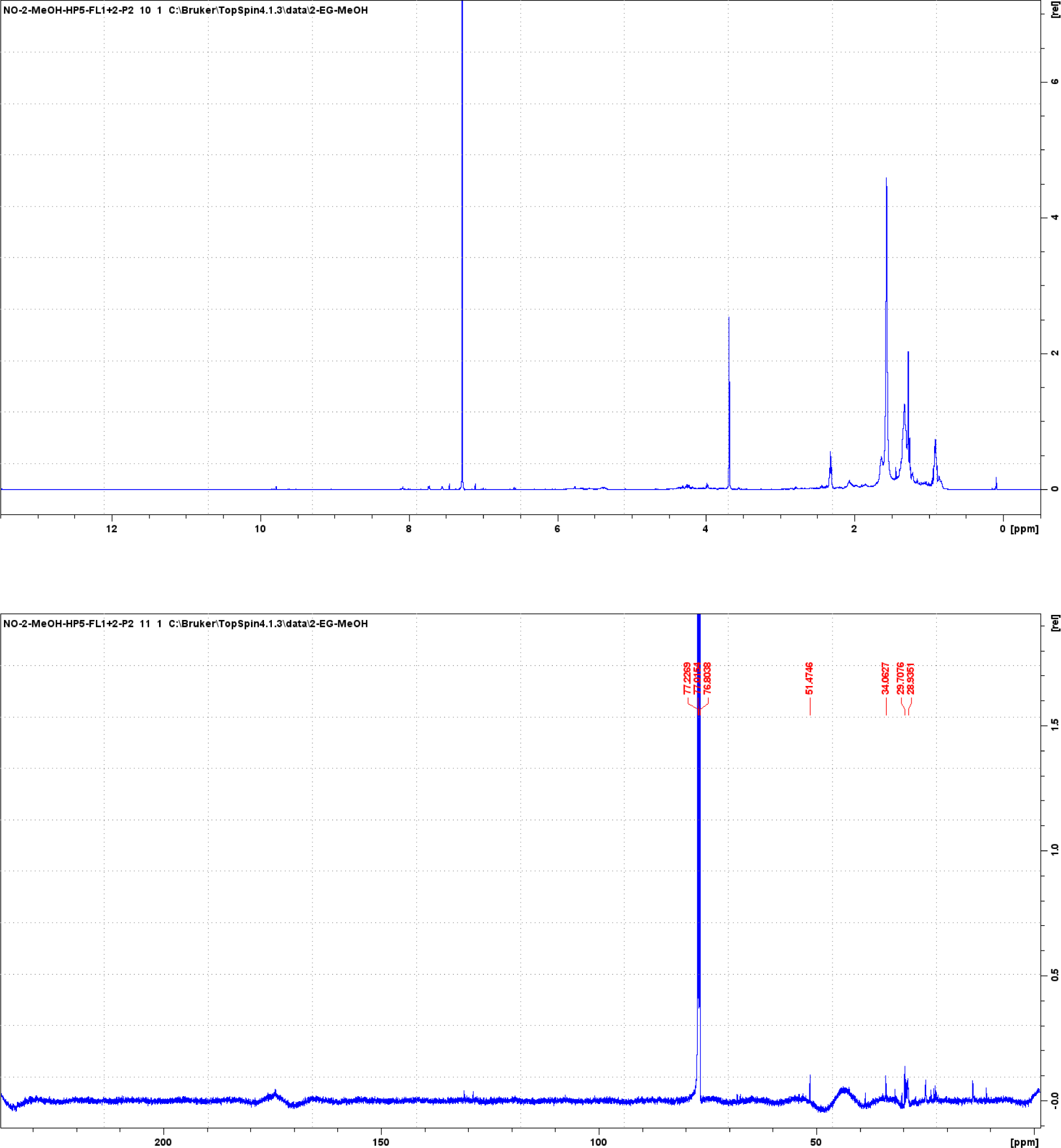
^1^H NMR (150 MHz) and ^13^C NMR (600 MHz) spectra of elaidic acid methyl ester (**8**) in CDCl_3_.

**FIG S3.**
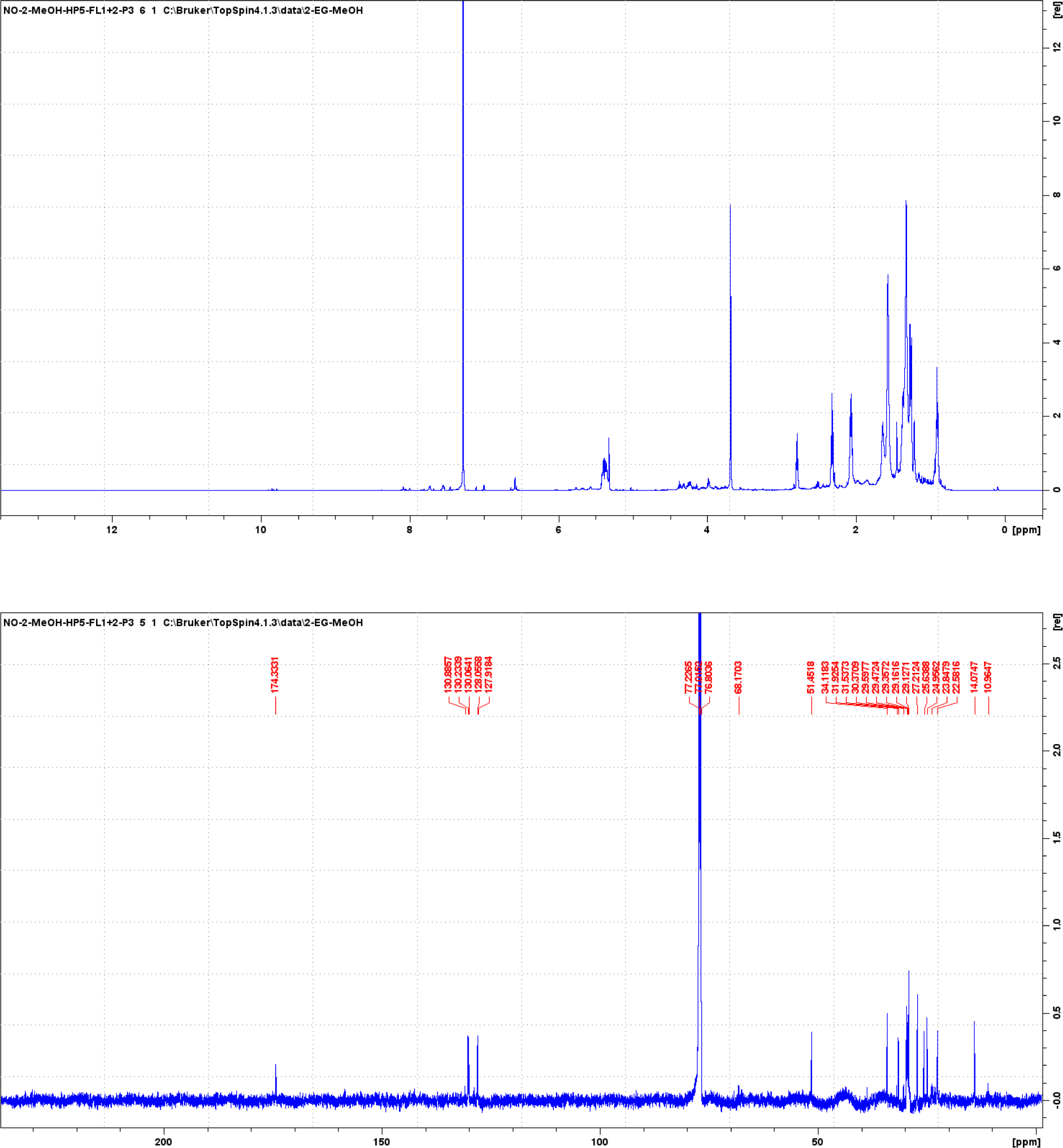
^1^H NMR (150 MHz) and ^13^C NMR (600 MHz) spectra of linoelaidic acid methyl ester (**9**) in CDCl_3_.

**FIG S4.**
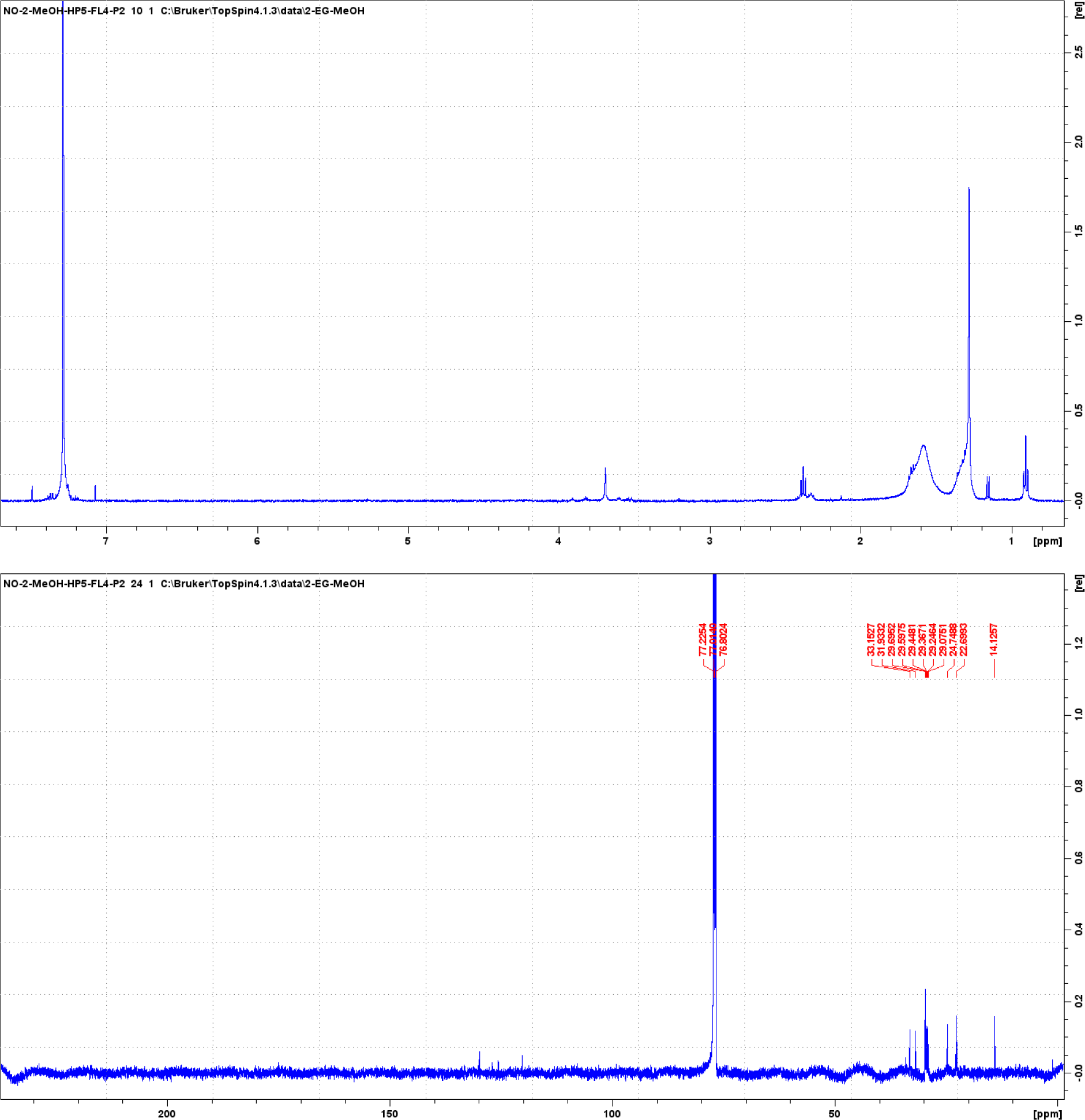
^1^H NMR (150 MHz) and ^13^C NMR (600 MHz) spectra of palmitic acid (**10**) in CDCl_3_.

**FIG S5.**
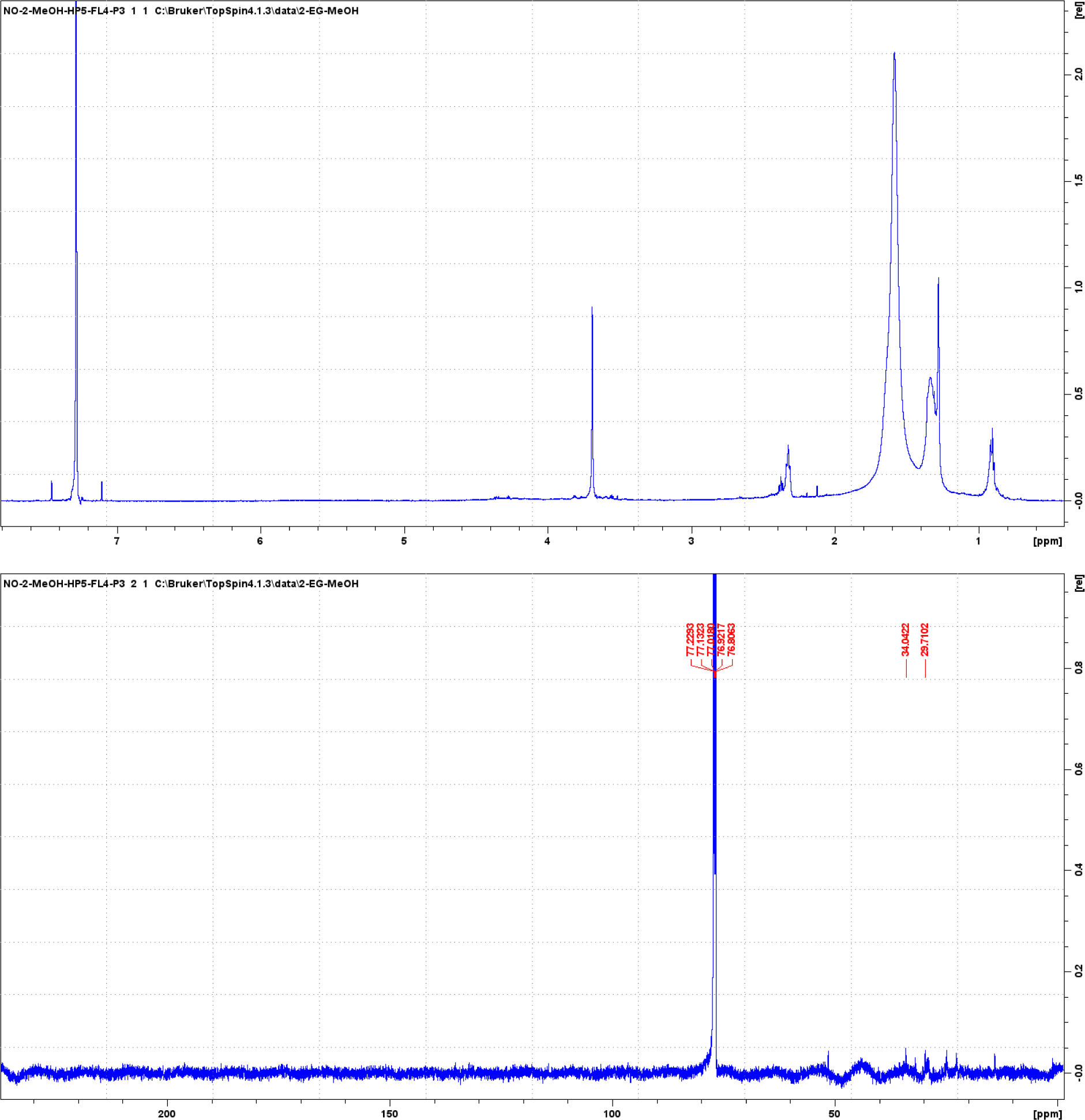
^1^H NMR (150 MHz) and ^13^C NMR (600 MHz) spectra of palmitic acid methyl ester (**11**) in CDCl_3_.

**FIG S6.**
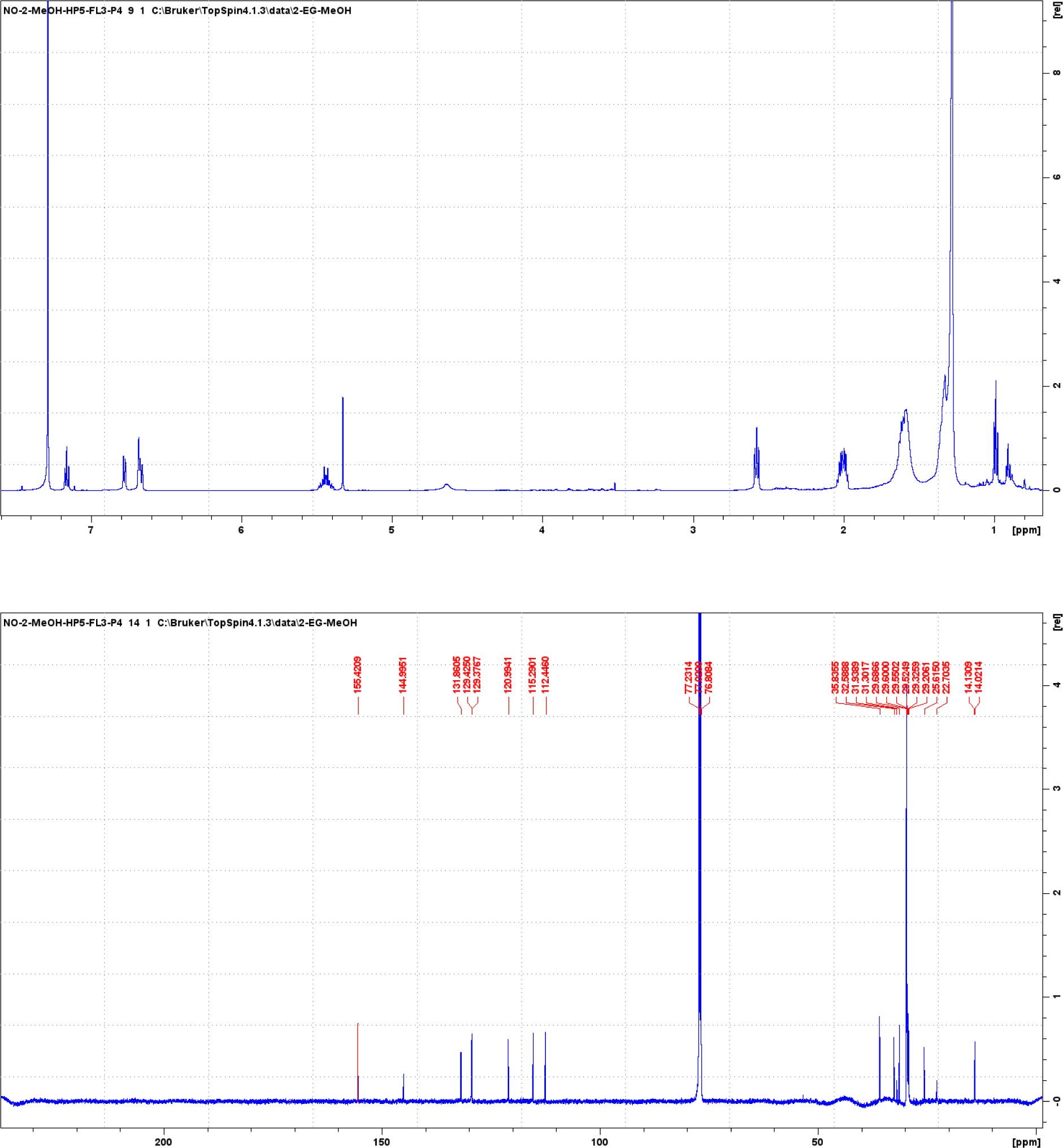
^1^H NMR (150 MHz) and ^13^C NMR (600 MHz) spectra of 3-[12(E)-pentadecenyl]phenol (**12**) in CDCl_3_.

**FIG S7.**
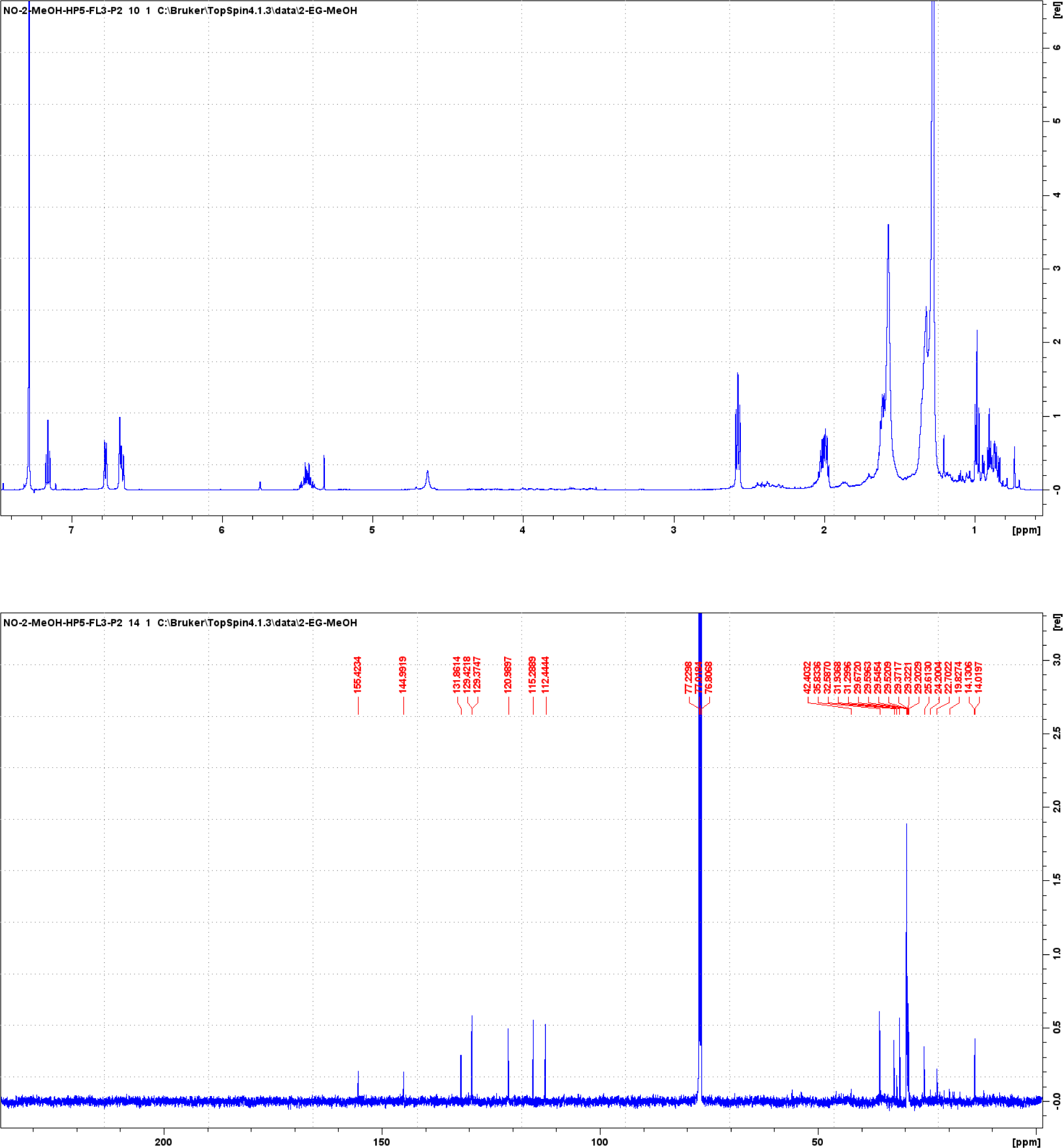
^1^H NMR (150 MHz) and ^13^C NMR (600 MHz) spectra of 3-[14(E)-heptadecenyl]phenol (**13**) in CDCl_3_.

**FIG S8.**
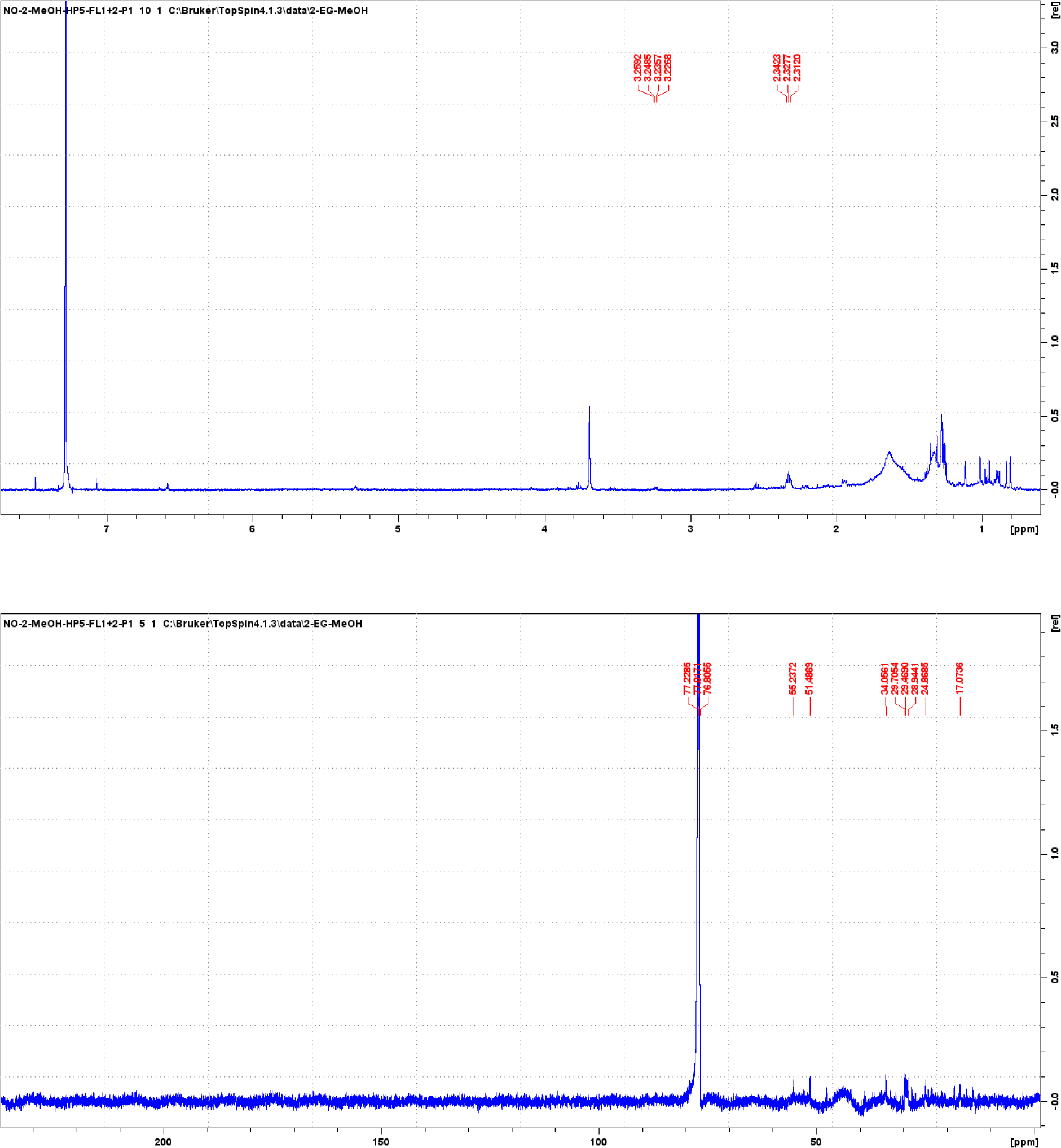
^1^H NMR (150 MHz) and ^13^C NMR (600 MHz) spectra of ursolic acid (**14**) in CDCl_3_.

**FIG S9.**
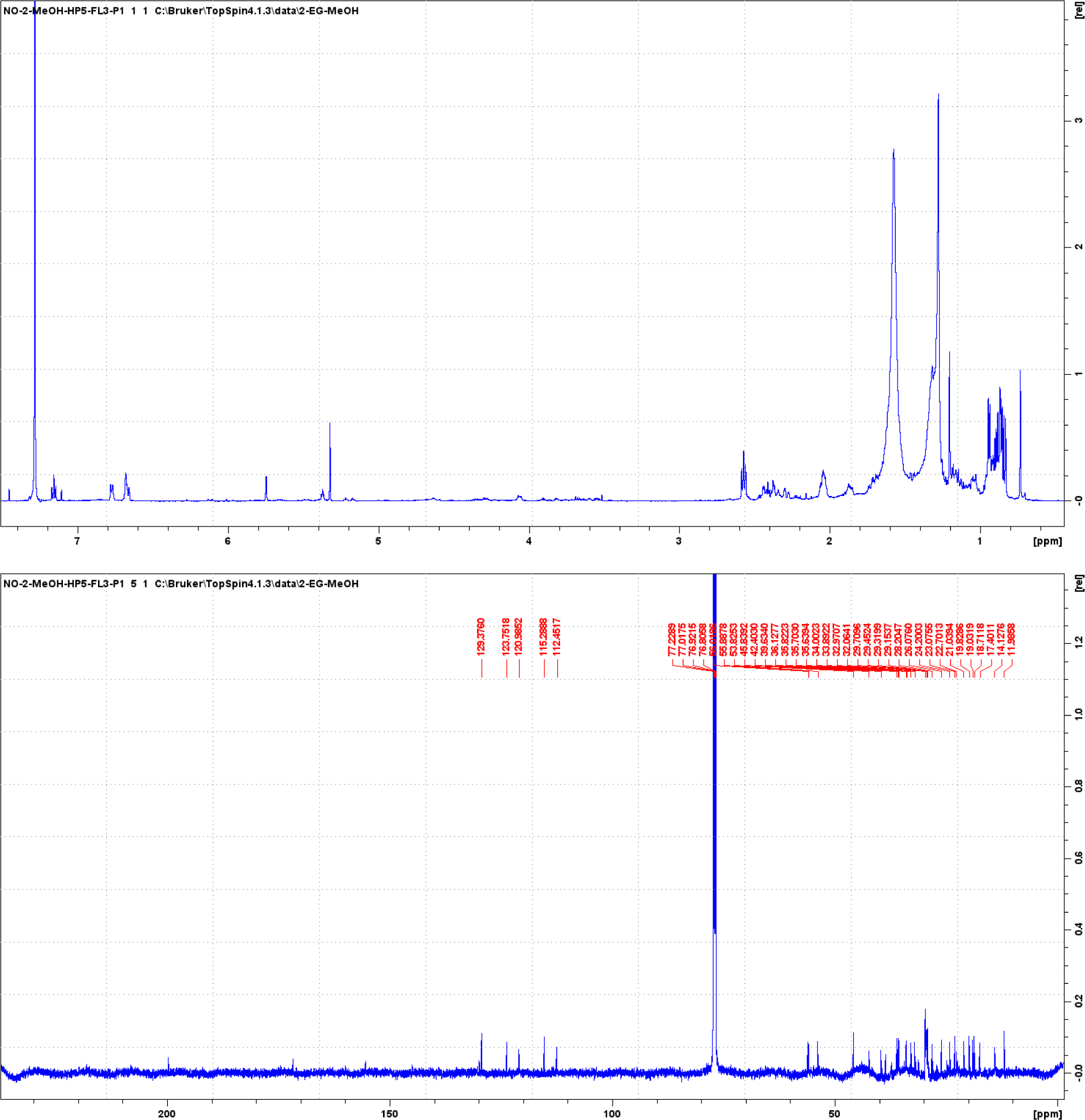
^1^H NMR (150 MHz) and ^13^C NMR (600 MHz) spectra of stigmasta-4,24(28)-dien-3-one (**15**) in CDCl_3_.

**FIG S10.** Image of plants harvested and subjected to the bio-guided pest control study.

**Table S1.**
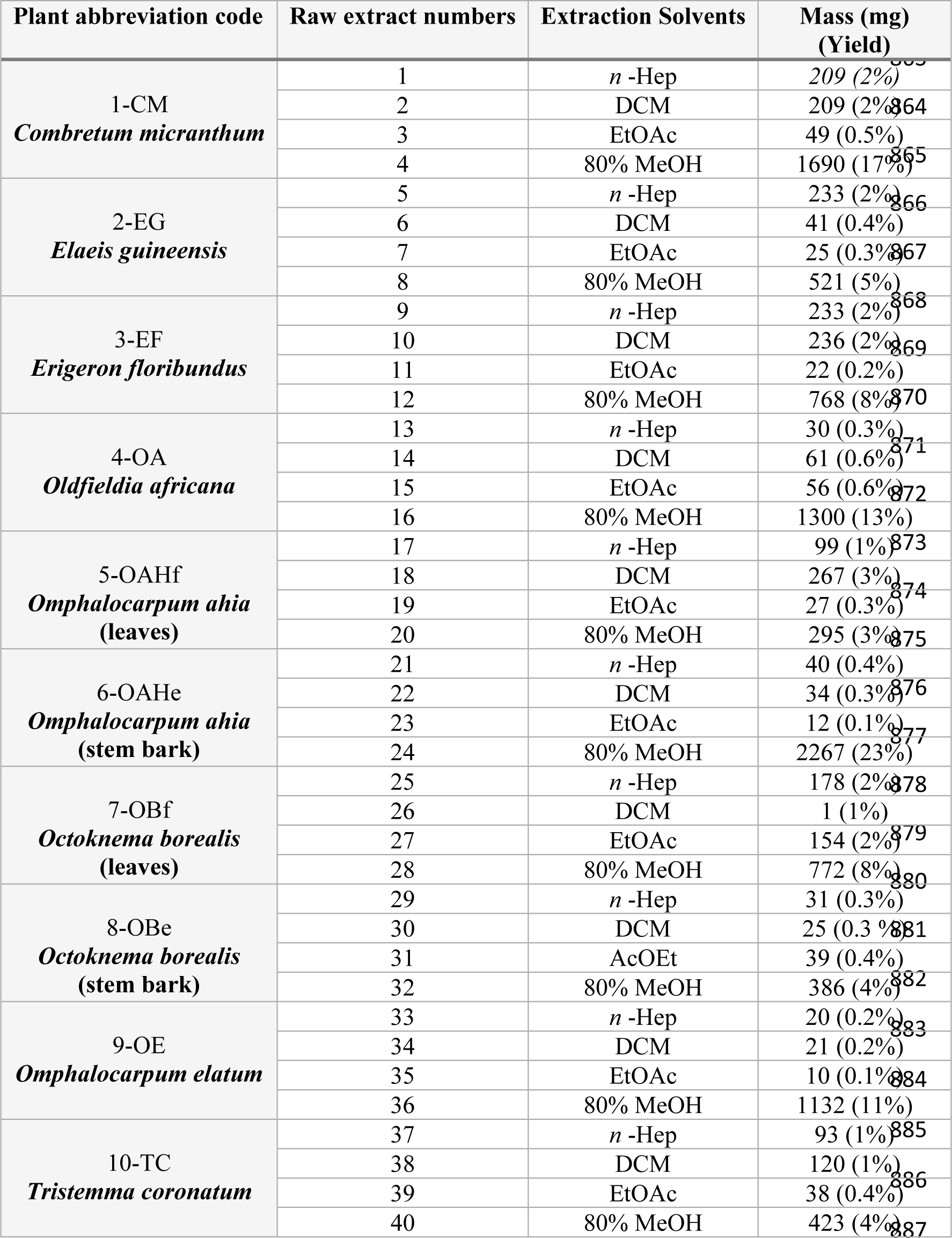
Numbering of the 40 extracts obtained after sequential extractions of increasing polarity, carried out on 10 g of each of the 10 ivorian plant species. Yields are given in brackets.

